# Multiple classes and isoforms of the RNA polymerase recycling motor protein HelD

**DOI:** 10.1101/2021.08.18.456904

**Authors:** Joachim S. Larsen, Michael Miller, Aaron J. Oakley, Nicholas E. Dixon, Peter J. Lewis

## Abstract

Efficient control of transcription is essential in all organisms. In bacteria, where DNA replication and transcription occur simultaneously, the replication machinery is at risk of colliding with highly abundant transcription complexes. This can be exacerbated by the fact that transcription complexes pause frequently. When pauses are long-lasting, the stalled complexes must be removed to prevent collisions with either another transcription complex or the replication machinery. HelD is a protein that represents a new class of ATP-dependent motor protein distantly related to helicases. It was first identified in the model Gram-positive bacterium *Bacillus subtilis* and is involved in removing and recycling stalled transcription complexes. To date, two classes of HelD have been identified: one in the low G+C and the other in the high G+C Gram-positive bacteria. In this work we have undertaken the first comprehensive investigation of the phylogenetic diversity of HelD proteins. We show that genes in certain bacterial classes have been inherited by horizontal gene transfer, many organisms contain multiple expressed isoforms of HelD, some of which are associated with antibiotic resistance, and that there is a third class of HelD protein found in Gram-negative bacteria. Therefore, HelD proteins represent an important new class of transcription factor associated with genome maintenance and antibiotic resistance that are conserved across the Eubacterial kingdom.

## INTRODUCTION

Transcription elongation is punctuated by pauses that serve important functions in permitting correct folding of structural RNA, efficient coupling of transcription and translation and ensuring efficient transcription termination at the correct site (Saba et al., 2019). Whilst most pausing events serve an important function, on occasion RNA polymerase (RNAP) is unable to restart transcription and must be removed from the DNA to prevent damaging collisions with the DNA replication machinery or other transcription complexes (Adelman and Lis, 2012, Gupta et al., 2013, Pomerantz and O’Donnell, 2008, Pomerantz and O’Donnell, 2010, Rocha, 2004). Several systems used to resolve stalled transcription complexes have been characterised; for example, Mfd has been shown to bind to stalled transcription complexes (either a stochastic pause during transcription of structured RNA or at a site of DNA damage), physically removing it from the DNA or restarting it *via* a RecG-like ATPase motor domain (Ragheb et al., 2021, Ghodke et al., 2020, Ho et al., 2018, Shi et al., 2020, Westblade et al., 2010, Kang et al., 2021, Le et al., 2018). In *B. subtilis* RNaseJ1 clears stalled RNAP using a torpedo mechanism (5’-3’ exonuclease activity followed by RNAP displacement) (Sikova et al., 2020), and in *Escherichia coli* the helicase protein RapA has been shown to be important in recycling RNAP (Liu et al., 2015). UvrD/PcrA in concert with Gre factors has been reported to act on RNAP stalled at a DNA lesion, binding to the complex and using the energy of ATP hydrolysis to backtrack away from the lesion to allow repair systems access to the damaged DNA (Epshtein et al., 2014, Hawkins et al., 2019), although it now appears that the role of these helicases is in preventing formation of, and resolving, R-loops (RNA-DNA hybrids) that can have a detrimental effect on DNA replication (Urrutia-Irazabal et al., 2021).

An additional system identified in Gram-positive bacteria required for recycling stalled transcription complexes involves the action of the motor protein HelD (Wiedermannova et al., 2014). The designation of HelD (also called helicase IV) was originally made for a protein identified in *E. coli* as a weakly processive 3’-5’ DNA helicase (Wood and Matson, 1987). To avoid confusion with the separate classes of HelD proteins that are the focus of this work, the *E. coli* protein will be referred to as helicase IV. Based on conserved sequence motifs Helicase IV is a superfamily 1 (SF1) helicase, related to housekeeping helicase UvrD/PcrA (Figure 1). The *B. subtilis* gene *yvgS* was assigned the name *helD* based on limited protein sequence conservation to helicase IV (Wiedermannova et al., 2014), although the proteins differed with respect to domain organisation (Koval et al., 2019, Wiedermannova et al., 2014)(Figure 1). Little functional, and no structural information is available for helicase IV, although a model generated by AlphaFold2 (Jumper et al., 2021) enables tentative comparison of UvrD/PcrA, helicase IV and *B. subtilis* HelD (Figure 1). Helicase IV and HelD show similarity with UvrD/PcrA around the well-defined 1A and 2A helicase domains (blue and orange, respectively, Figure 1A), but not in other structural motifs associated with helicase activity (UvrD/PcrA domains 1B and 2B). Both helicase IV and HelD have N-terminal domains not present in UvrD/PcrA helicases, and helicase IV has a putative 1B domain which may account for its reported helicase activity, whilst in the equivalent 1B domain position HelD contains unrelated sequence that folds into a novel clamp-arm (CA) structure important in transcription recycling (Newing et al., 2020, Wiedermannova et al., 2014). Whilst UvrD/PcrA and helicase IV have helicase activity, HelD shows none suggesting it has evolved from an SF1-type helicase into a transcription recycling factor that utilises the energy from ATP hydrolysis catalysed by its helicase motifs for its transcription-related activity.

**Figure 1.**
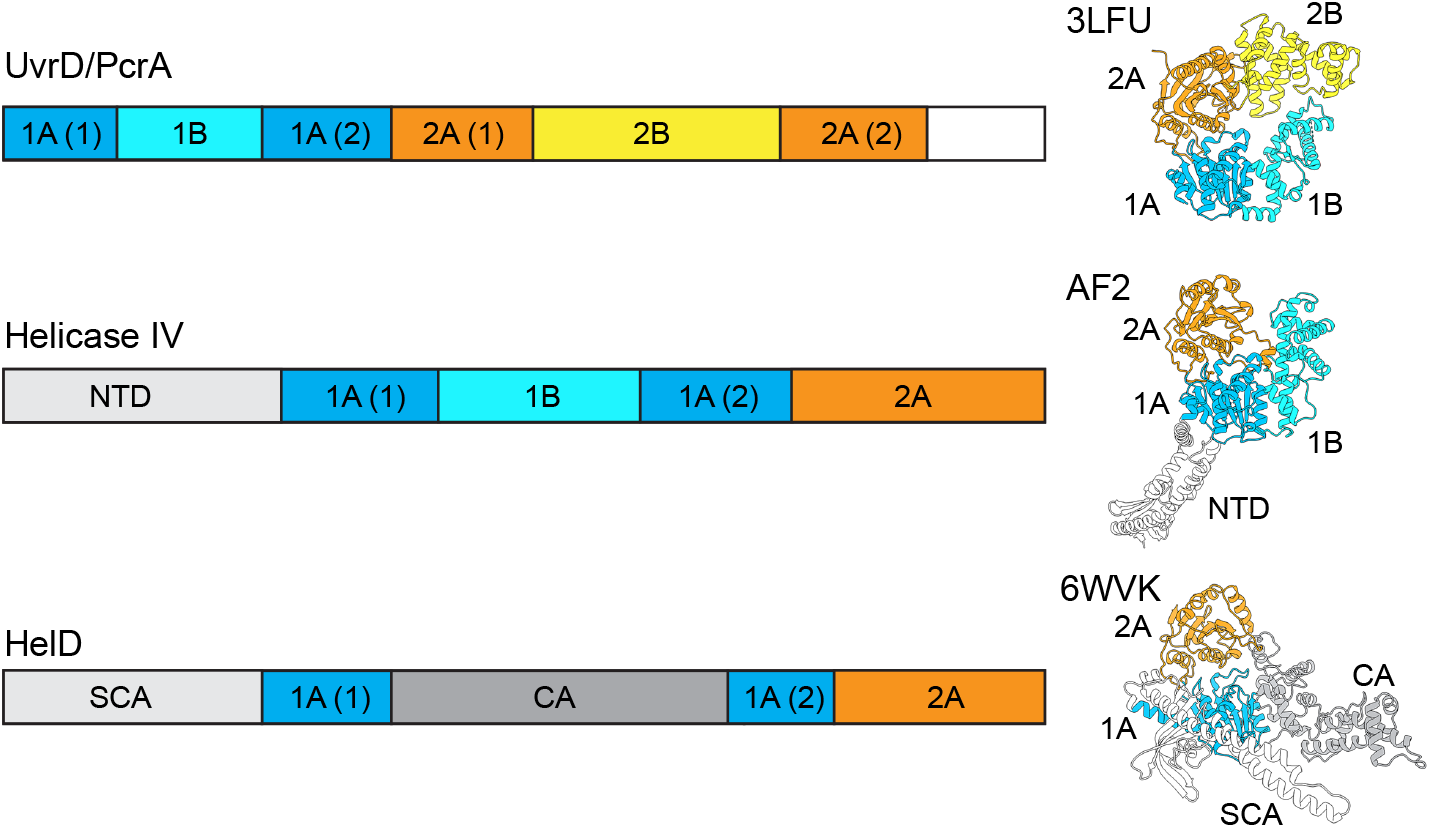
Relationship between UvrD/PcrA and helicase IV/HelD proteins. Left side shows linear representations of the domain organisation of superfamily 1 (SF1) helicase UvrD/PcrA (top), *Escherichia coli* helicase IV (middle) and *B. subtilis* HelD (bottom). Right hand side shows structures, aligned *via* their 1A and 2A domains, with domains coloured corresponding to the left panels. Top, UvrD (PDB ID 3LFU); middle, helicase IV (AlphaFold2 model, AF2); bottom, HelD (taken from RNAP-HelD complex PDB ID 6WVK). 1A, B, 2A and 2B refer to conserved SF1 helicase domains. NTD, SCA and CA refer to the AlphaFold2 modelled N-terminal domain of helicase IV and the secondary channel arm and clamp arm of HelD, respectively.

Studies on HelD from low G+C (*Bacillus subtilis*) and high G+C (*Mycobacterium smegmatis*) Gram-positives revealed that there are two distinct classes of enzyme, confirmed by phylogenetic and structural analyses (Kouba et al., 2020, Newing et al., 2020, Pei et al., 2020). Class I HelD was described from *B. subtilis*, whilst the structurally distinct Class II enzyme was identified in *M. smegmatis* (Kouba et al., 2020, Newing et al., 2020, Pei et al., 2020). Class I and II HelDs have similar motor domains but differ in the structure of their arms and the mechanism by which these arms perform the mechanical activity of removing nucleic acids and recycling RNAP (Kouba et al., 2020, Newing et al., 2020, Pei et al., 2020).

The recent structures of HelD from *B. subtilis* and *M. smegmatis* bound to core RNAP (α_2_ββ’ω) (Kouba et al., 2020, Newing et al., 2020) are shown in Figure 2A and B, along with the Class I *B. subtilis* (Figure 2C) and Class II *M. smegmatis* (Figure 2D) enzymes. HelD has an unusual mode of action dependent on two arms (CA and SCA, Figure 2C and D) attached to the central UvrD-like ATPase motor domain (Head and Torso, Figure 2C and D), in which nucleic acids are pushed out of the active site whilst the DNA binding clamp and RNA exit channels are simultaneously opened, leading to the release of the stalled RNAP (Newing et al., 2020). This recycling activity is powered by ATP hydrolysis and the mechanical action of the two arms that flank the motor domain. In the Class I HelD, the long SCA (Figure 2A and C) is able to physically remove nucleic acids from the active site (dotted circle in Figure 2A), whereas in the Class II HelD the SCA is too short, and instead nucleic acid removal is performed by a CA insert called the PCh-loop (Figure 2B and D) (Kouba et al., 2020, Newing et al., 2020). Recent reports also suggest that some Class II HelDs (from *M. abscessus* and *Streptomyces venezuelae*) are able to confer rifampicin resistance through removal of rifampicin by the PCh-loop (Hurst-Hess et al., 2021, Surette et al., 2021).

**Figure 2.**
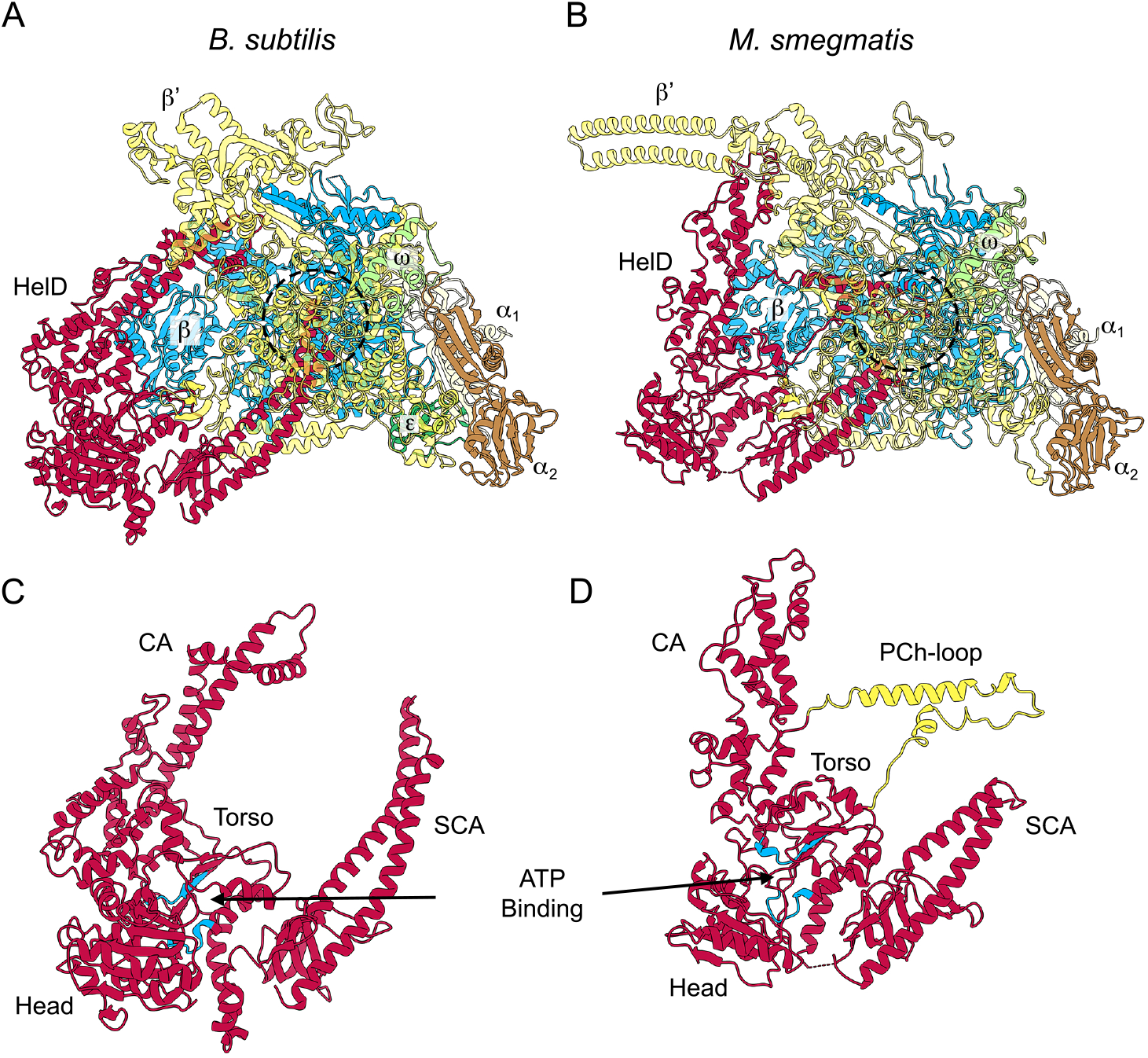
The two known structural classes of HelD. Panel A shows the structure of the *B. subtilis* RNAP-Class I HelD complex (PDB ID 6WVK). Panel B shows the *M. smegmatis* RNAP-Class II HelD complex (PDB ID 6YYS; state II). RNAP subunits and HelDs are coloured identically in both panels with the transparency of the β’ subunit set at 50% so that HelD structures adjacent to the RNAP active site region (dashed circles) can be more easily visualised. Panels C and D show HelD structures from Panels A and B, respectively, with the ATP binding site coloured in blue and the PCh-loop from *M. smegmatis* HelD coloured in yellow (see text for details).

In this work, we take advantage of the recent structural information to compile a detailed phylogenetic analysis of HelD showing that many organisms contain more than one (up to 5) different versions of HelD, that the genes encoding these enzymes are all expressed, that HelD is likely to have been acquired by horizontal gene transfer in Gram-negative *Bacteroides* and Gram-positive *Coriobacteria* and *Acidimicrobiia*, and that there is a third Class of HelD found in the Gram-negative *Deltaproteobacteria*.

## RESULTS AND DISCUSSION

### Distribution and phylogeny of HelD

Searching for HelD-like sequences using the conserved domain architecture retrieval tool (CDART; NCBI) portal identified >13,000 hits. Additional searches using NCBI BLASTP suggest that there are substantially more sequences in the database, but many of these are from incomplete genomes and/or metagenomic sequencing projects, making systematic identification and classification of sequences unfeasible, particularly in cases where an organism carries more than one *helD* gene (see below). Nevertheless, it is clear that HelD is widely distributed in the eubacteria, especially in the *Firmicutes* and *Actinobacteria* phyla of the Gram-positive eubacterial domain. To date, we have not detected HelD-like sequences in *Archaea* or *Eucarya*. Previously, Newing et al. (Newing et al., 2020) showed that HelD sequences fall into two classes, which was confirmed at the structural and functional level in comparing HelD proteins from the *Firmicutes* and *Actinobacteria* (Kouba et al., 2020, Newing et al., 2020, Pei et al., 2020). Using a wider range of carefully curated sequences from complete genomes identified from the initial CDART search, an unrooted phylogenetic tree was constructed to enable a more detailed understanding of HelD distribution and phylogeny which was compared against the RNAP RpoB (β) subunit (Figure 3).

**Figure 3.**
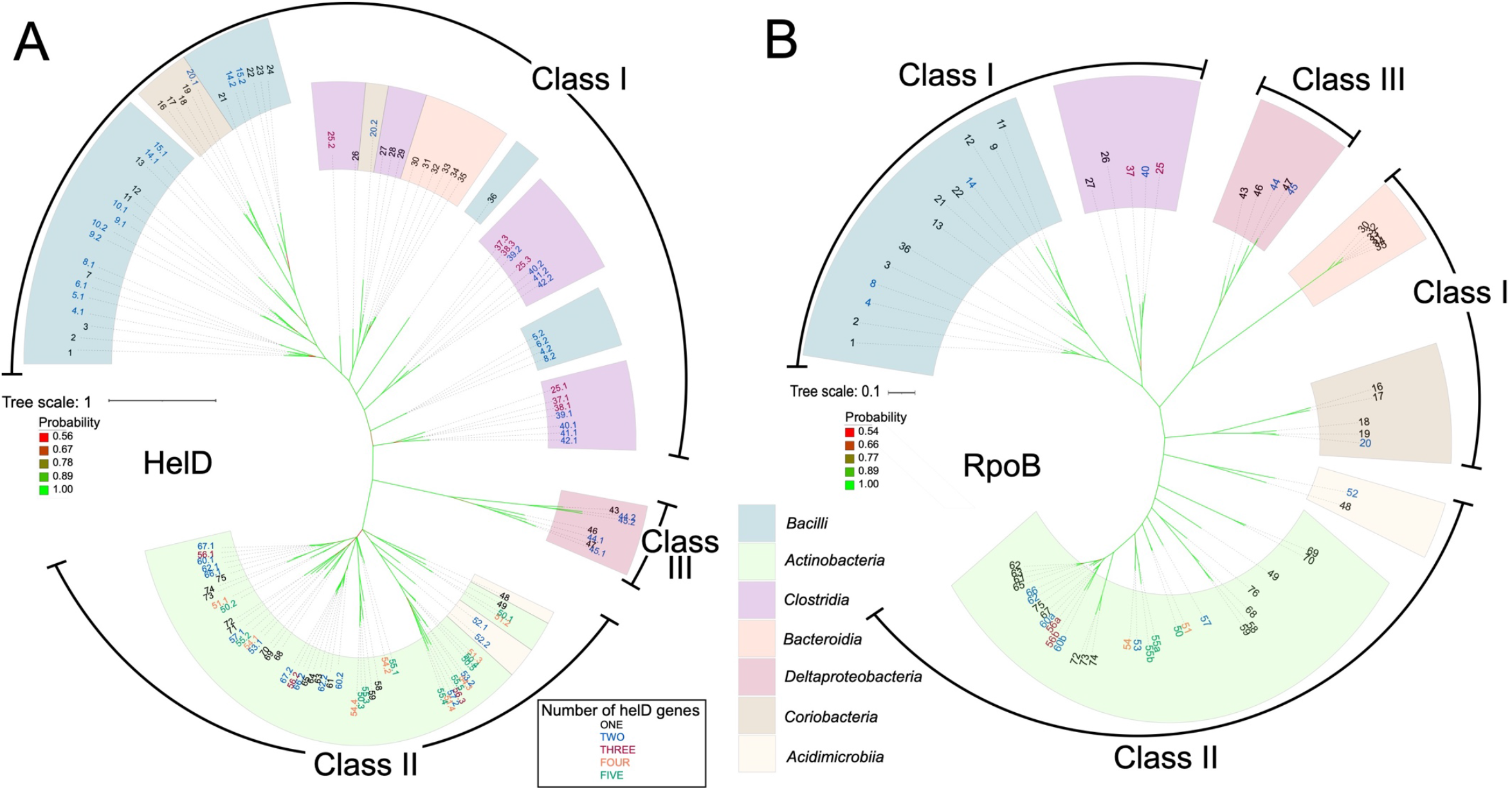
Unrooted phlyogenetic trees of HelD (A) and RpoB (B) sequences constructed by Bayesean analysis. Tree scale representing amino acid substitutions per site, and bootstrap probability values (red least, to green most, probable) are on the left. The HelD class into which sequences fall is indicated in the outer circles as Class I, -II and -III. Coloured arcs indicate the bacterial classes into which the HelD sequences fall; teal, *Firmicutes*; pale green, *Actinobacteria*; purple, *Clostridia*; orange, *Bacteroidia*; red, *Deltaproteobacteria*; brown, *Coriobacteria*; pale yellow, *Acidimicrobilia*. Individual organisms and HelD sequences are numbered (largest to smallest) and colour coded starting clockwise from *Bacillus subtilis*. Organism numbers with one HelD are numbered in black; two, blue; three, red; four, orange; five, green and are listed as follows with gene identifiers and protein length (aa) in brackets: **1** *Bacillus subtilis* 168 (BSU_33450, 774aa). **2** *Bacillus licheniformis* ATCC 14580 (bli_00699, 776aa). **3** *Bacillus megaterium DSM 319* (BMD_3869, 772aa). **4** *Bacillus cereus* ATCC10987 (#1 BCE_3516, 768 aa; #2 BCE_2839, 689 aa). **5** *Bacillus anthracis AMES* (#1 BA_1040, 776 aa; #2 BA_2814, 689 aa). **6** *Bacillus cereus* AH187 (#1 BCAH187_A1206, 777 aa; #2 BCAH187_A2861, 689 aa). **7** *Bacillus cereus* ATCC14579 (BC_1041, 777 aa). **8** *Bacillus thuringiensis* Bt407 (#1 btg_c11000, 778aa; #2 btg_c29280, 691aa). **9** *Lactobacillus plantarum* WCFS1 (#1 lpl_0432, 769aa; #2 lpl_0910, 768aa). **10** *Lactobacillus rhamnosus* GG (#1 lrh_01975, 763aa; #2 lrh_02619, 762aa). **11** *Leuconostoc lactis* WiKim40 (llf_04535, 788aa). **12** *Lactobacillus acidophilus* NCFM (lac_1676, 687aa). **13** *Carnobacterium inhibens* subsp. Gilchinskyi WN1359 (caw_09345, 800aa). **14** *Enterococcus faecium* Aus0004 (#1 EFAU004_01304, 759 aa; #2 EFAU004_00387, 711 aa). **15** *Enterococcus faecium* DO (#1 HMPREF0351_10989, 759 aa; #2 HMPREF0351_10397, 711 aa). **16** *Olsenella uli* DSM 7084 (OLS_0501, 731aa). **17** *Atopobium parvulum* DSM 20469 (Apar_0360, 736aa). **18** *Slackia heliotrinireducens* DSM 20476: (Shel_05840 (698aa). **19** *Eggerthella lenta* DSM 2243(Elen_2835, 716aa). **20** *Adlercreutzia equolifaciens* DSM 19450 (#1 AEQU_1689, 761aa; #2 AEQU_0484, 733aa). **21** *Vagococcus teuberi* (vte_03205, 717aa). **22** *Enterococcus faecalis* V583 (EF_0933, 732 aa). **23** *Enterococcus faecalis* DENG1 (DENG_00988, 732 aa). **24** *Enterococcus faecalis* OG1RF (OG1RF_10660, 740 aa). **25** *Clostridium beijerinckii* NCIMB 8052 (#1 cbe_2947, 755aa; #2 cbe_2724, 745aa; #3 cbe_4782, 724aa). **26** *Epulopiscium sp*. N.t. morphotype B (EPU_RS03295, 735aa). **27** *Clostridioides difficile* 630 (CD630_04550, 704 aa). **28** *Clostridioides difficile* RM20291 (CDR20291_0396, 704 aa). **29** *Clostridioides difficile* CD196 (CD196_0410, 704 aa). **30** *Bacteroides vulgatus* ATCC 8482 (BVU_3010 (671aa). **31** *Bacteroides caccae* ATCC 43185 (CGC64_00555, 683aa). **32** *Bacteroides cellulosilyticus* WH2 (BcelWH2_01491, 693aa). **33** *Bacteroides thetaiotaomicron* VPI-5482 (BT_1890, 686aa). **34** *Bacteroides ovatus* ATCC 8483 (Bovatus_02598 (687aa). **35** *Bacteroides xylanisolvens* XB1A (BXY_17560, 687aa). **36** *Staphylococcus delphini* NCTC12225 (sdp_01978, 681aa). **37** *Clostridium botulinuim A* ATCC3502 (#1 CBO_2904, 763 aa; #3 CBO_3341, 709 aa). **38** *Clostridium botulinuim A* ATCC19377 (#1 CLB_2867, 763 aa; #3 CLB_3399, 709 aa). **39** *Clostridium botulinuim B1 Okra* (#1 CLD_1639, 763 aa; #2 CLD_1179, 709 aa). **40** *Clostridium perfringens* 13 (#1 CPE_1619, 763 aa; #2 CPE_0599, 706 aa). **41** *Clostridium perfringens* ATCC13124 (#1 CPF_1872, 763 aa; #2 CPF_0580, 706 aa). **42** *Clostridium perfringens* SM101 (#1 CPR_1591, 763 aa; #2 CPR_0566 706 aa). **43** *Myxococcus xanthus* DK 1622 (MXAN_5482, 706aa). **44** *Sandaracinus amylolyticus* DSM 53668 (#1 DB32_004372, 872aa; #2 DB32_003397, 691aa). **45** *Minicystis rosea* DSM 2400 (#1 A7982_09686, 743aa; #2 A7982_06548, 703aa). **46** *Haliangium ochraceum* DSM 14365 (Hoch_0025, 852aa). **47** *Sorangium cellulosum* So157-2 (SCE1572_03860, 747aa). **48** *Acidobacterium ferrooxidans* (Afer_1829, 706aa). **49** *Cutibacterium acnes* KPA171202 (PPA0733, 753aa). **50** *Streptomyces venezuelae* (#1 SVEN_2719, 779aa; #2 SVEN_5092, 747aa; #3 SVEN_6029, 722aa; #4 SVEN_4127, 675aa; #5 SVEN_3939<u>; 665aa</u>). **51** *Streptomyces coelicolor* A3(2) (#1 SCO5439, 755 aa; #2 SCO2952, 744 aa; #3 SCO4316, 681 aa; #4 SCO4195, 680 aa). **52** *Ilumatobacter coccineus* (#1 aym_09360, 715aa; #2 aym_20540, 654aa). **53** *Frankia casuarinae* Ccl3 (#1 fra_0952, 829aa; #2 fra_2397, 727aa). **54** *Frankia alni* ACN14a (#1 fal_1589, 939aa; #2 fal_4723, 877aa; #3 fal_3805; 866aa; #4 fal_4811, 751aa). **55** *Nonomuraea sp*. ATCC55076 (#1 NOA_23645, 772 aa; #2 NOA_16240, 762 aa; #3 NOA_42280, 715 aa; #4 NOA_08745, 660 aa; #5 NOA_48960, 655 aa). **56** *Nocardia brasiliensis* O31_020410 (#1 nbr_012985, 776aa; #2 nbr_020410, 731aa; #3 nbr: O3I_005870, 699aa). **57** *Kineococcus radiotolerans* SRS30216 (#1 kra_3607, 759aa; #2 kra_0164, 684aa). **58** *Microbacterium sp*. PAMC 28756 (mip_00070, 717aa). **59** *Mirobacterium hominis* SJTG1 (mhos_01135, 744aa). **60** *Nocardia farcinica* IFM10152 (#1 NFA_19060, 765aa; #2 NFA_44160, 726aa). **61** *Mycobacterium smegmatis* MC2 155 (MSMEG_2174, 736aa). **62** *Rhodococcus sp*. 008 (#1 rhod_26990, 760aa; #2 rhod_09075, 731aa). **63** *Mycobacterium sp*. JS623 (Mycsm_03949, 732aa). **64** *Mycolicibacterium phlei* (MPHL_03003, 726aa). **65** *Mycobacteroides abscessus* ATCC 19977 (MAB_3189c, 753aa). **66** *Rhodococcus equi* 103S (#1 REQ_25070, 759aa; #2 REQ_15310, 739aa). **67** *Nocardia asteroides* NCTC11293 (#1 nad_03000, 753; #2 nad_04408, 735aa). **68** *Leifsonia xyxli* subsp. Xyli CTCB07 (Lxx_20770, 787aa). **69** *Bifidobacterium longum* NCC2705 (BLO_1314, 759aa). **70** *Bifodobacterium bifidum* PRL2010 (bbp_0546, 759aa). **71** *Brevibacterium linens* BS258 (bly_10570, 743aa). **72** *Brevibacterium flavum* ZL-1 (bfv_07580, 755aa). **73** *Corynebacterium glutamicum* ATCC13031 (CG_1555, 755aa). **74** *Corynebacterium diptheriae* NTCC13129 (DIP_1156, 770aa). **75** *Rhodococcus rhodochrous* NCTC10210 (rrt_02795, 772aa). *Nonomuraea sp*. ATCC55076 (55), *Nocardia brasiliensis* O31_020410 (56) and *Nocardia farcinica* IFM10152 (60) contain two copies of the *rpoB* gene (numbered x.a and x.b in panel B). Copy 1 is the housekeeping *rpoB* and copy 2 is a rifamipicin-resistant *rpoB* expressed during antibiotic production in those organisms.

Four features are clear from this tree (Figure 3A): 1. HelD is also present in Gram-negative bacteria; 2. A third class of HelD is present in the *Deltaproteobacteria*; 3. In some organisms HelD has been ancestrally acquired by horizontal gene transfer; 4. Many organisms contain more than one *helD* gene, with the *Firmicutes, Clostridia, Acidimicrobiia*, and *Deltaproteobacteria* having up to three, and the *Actinobacteria* up to five.

Overall, the tree contains three major branches: Class I HelD sequences originating mainly from the low G+C Gram-positives and *Bacteriodia*, Class II HelD sequences from the high G+C Gram-positives, and a novel Class III identified in *Deltaproteobacteria*. Interestingly, the HelD sequences from the Actinobacterial *Coriobacteria* class, typified by *Olsenella uli* that is associated with gingivitis, are all located to the Class I branch of the tree (numbers 16–20; Figure 3). Branch divergence and clustering of sequences to regions of the tree comprising *Lactobacilli* (numbers 14, 15, 21–24; Figure 3) and *Clostridia* (numbers 25–29; Figure 3) indicate that an ancestral *Coriobacteria* likely acquired *helD* genes by horizontal gene transfer from these organisms (Figure S1). That *Coriobacteria* are isolated from the gingival crevice, gastrointestinal and genital tracts (Clavel, 2014) is consistent with this proposition. The length of the branches suggests this horizontal transfer event occurred long ago but after the evolution of the mammalian hosts that provide environments with co-localised *Lactobacilli*, and that *helD* genes have been stably inherited and co-evolved within the *Coriobacteria*. In addition to the *helD* gene from *Adlercreutzia equolifaciens* DSM 19450 (AEQU_1689, number 20.1; Figure 3) that clusters with those of the other *Coriobacteria, A. equolifaciens* contains a second *helD* gene (AEQU_0484, number 20.2; Figure 3) that clusters with *Clostridia*, suggesting it may have been acquired through a separate horizontal gene transfer event rather than through duplication and evolution of a gene inherited by a single acquisition event (Figure S1). The fact that *Lactobacilli, Clostridia*, and *Aldercreutzia* all inhabit the gastrointestinal tract make this a reasonable hypothesis. There is also some evidence that Class II HelD sequences have been acquired by horizontal gene transfer from the *Actinobacteria* to the *Acidimicrobiia* (numbers 48, 52.1 and 52.2; Figures 3 and S2). The *Acidimicrobiia* are a recently described class, exemplified by *Acidobacterium ferrooxidans* (number 48; Figure 3) that have been isolated from diverse, but generally acidic and hostile environments, and tend to grow slowly which may account for the paucity of information and diversity of species currently available. At least one species of the *Acidimicrobiia, Ilumatobacter coccineus* (number 52, Figure 3) contains multiple copies of *helD*.

Comparison of the phylogenetic tree of the RNA polymerase β subunit RpoB with the HelD tree supports this assumption that *helD* genes in the *Coriobacteria* and *Acidimicrobiia* have been acquired by horizontal gene transfer from *Firmicutes/Clostridia/Actinobacteria* that share the same ecological niches (Figures 3A and B). Acquisition of *helD* genes by horizontal gene transfer in the *Bacteroidia* is described below.

### Acquisition of *helD* in Gram-negative *Bacteroides*

HelD sequences were also identified in the phylum of Gram-negative bacteria, *Bacteroides*. Phylogenetically, these clustered close to HelD sequences from *Clostridioides difficile* (Figures 3A and S3; sequences 27–29 *C. difficile*, 30–35 *Bacteroides*). Extended analysis indicated that HelD sequences from *Bacteroides* and *Parabacteroides* (family *Porphyromonadaceae*) clustered closest to those from *Firmicutes* that are strict gut anaerobes from the order *Clostridiales* (Figure S4). These bacteria were from cluster IV (*Ruminoccoaceae*) and XIVa (*Lachnospiraceae*) that are abundant gut microbes associated with many aspects of good health, and the cluster XI gut pathogen *C. difficile* (Lopetuso et al., 2013, Lozupone et al., 2012, Milani et al., 2017). Since the *Bacteroides* and *Parabacteroides* are also abundant obligate gut anaerobes, this clustering suggested that *helD* was horizontally transferred from an anaerobic gut *Firmicute*, most likely from the order *Clostridiales* (Figure S3). Analysis of the genome context of *helD* genes indicated they were not (or are no longer) located in mobile genetic elements, with the exception of *B. thetaiaotamicron*, and along with their widespread distribution in *Bacteroides*/*Parabacteroides* suggests *helD* genes have been retained over a significant time period, indicating they serve a useful cellular function. The fact that HelD sequences identified in *Bacterioides* cluster with Class I sequences from the low G+C Gram-positive bacteria rather than forming a separate Class, as seen with HelD from the *Deltaproteobacteria* (see below), further supports the idea that this group acquired *helD* genes by horizontal gene transfer due to sharing a similar environmental niche to anaerobic gut *Clostridiales*.

### A novel HelD class in Gram-negative bacteria

The analysis presented in this work also shows that there is a third class of HelD proteins encoded by the *Deltaproteobacteria* (Class III, Figure 3 and 4; see below). Newing *et al*. (Newing et al., 2020) identified Class I and II HelD proteins based on the conservation of twelve sequence motifs. These motifs (labelled I-XII, Figure S5) are all conserved in Class III proteins (exemplified by *Myxococcus xanthus* HelD), despite the low overall levels of sequence similarity found in HelD proteins (Newing et al., 2020). A model of *M. xanthus* HelD was also generated from an unbiased screen of the protein structure database (Figure 4; see Materials and Methods). As seen with Class I and II proteins, there is a HelD-specific N-terminal domain of ∼50–150 amino acids that has a long antiparallel α-helical structure (secondary channel arm, SCA, Figure 4B) that is required to anchor HelD in the secondary channel of its cognate RNAP (Kouba et al., 2020, Newing et al., 2020, Pei et al., 2020), and the 1A helicase domain is split by the insertion of an arm-like structure (clamp arm, CA, Figures 4B and S5) that is used to bind within the primary channel of RNAP, forcing it open to aid the release of bound nucleic acids (Kouba et al., 2020, Newing et al., 2020, Pei et al., 2020).

**Figure 4.**
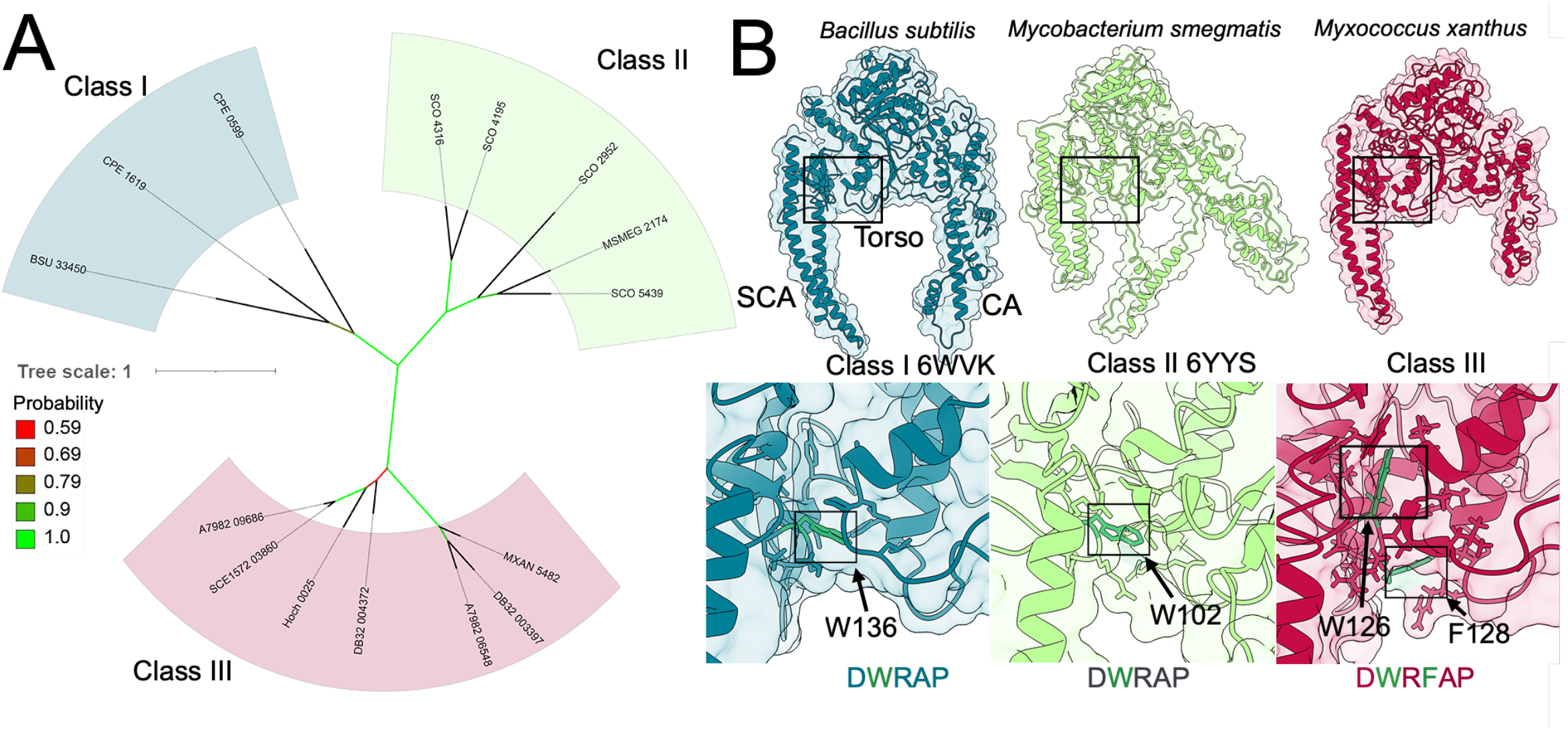
Three classes of HelD. Panel A shows a focused unrooted phylogenetic tree constructed using HelD sequences, with numbers (#) as used in Figure 1A: *B. subtilis* 168, BSU [#1]; *C. perfringens* 13, CPE [#40]; *S. coelicolor* A3(2), SCO [#51]; *M. smegmatis* MC2 155, MSMEG [#61], and *Deltaproteobacterial* sequences from *M. xanthus* DK 1622, MXAN [#43]; *S. amylolyticus* DSM 53668, DB32 [#44]; *M. rosea* DSM 2400, A7982 [#45]; *H. ochraceum* DSM 14365, Hoch [#46]; *S. cellulosum* So157-2, SCE1572 [#47]. Tree scale representing amino acid substitutions per site, and bootstrap values are shown on the left. Colouring of bacterial classes is the same as in Figure 1. Panel B shows structures (ribbons and transparent surface representations) of whole HelD (top) and Trp-cage regions (bottom) of Class I (*B. subtilis* PDB ID 6WVK), Class II (*M. smegmatis* PDB ID 6YYS) and Class III (*M. xanthus*, homology model) using the same colour scheme for bacterial classes as in Figures 1 and 2A. Conserved Trp (all classes) and additional amino acid (Class III) are shown as green sticks, with other amino acids that form the cage shown in the appropriate colour for their class.

An absolutely conserved DWR (Asp-Trp-Arg) sequence motif was identified in the unique N-terminal domain of all HelD sequences, and determination of the structures of HelD showed that the conserved Trp residue resides within a hydrophobic pocket called the Trp-cage, important in stabilising the interaction between the N-terminal domain wedged deep into the secondary channel of RNAP and the helicase 1A domain (Newing et al., 2020). In most HelD sequences identified to date, the DWR motif is extended to DWR[A/S]P, but in *Deltaproteobacterial* HelDs there is an additional amino acid inserted in this motif following the R residue, i.e., DWR<u>X</u>[A/S]P, which is a key defining feature of a Class III HelD (Figure S5). This additional amino acid does not appear to be highly conserved, the motif being DWR<u>F</u>AP in *M. xanthus*, DWR<u>N</u>AP in *Haliangum ocraceum*, and DWR<u>H</u>AP in *Sorangium cellulosum*, with H or N appearing to be most common. Modelling suggests this amino acid is located on a loop with its side chain in an additional pocket that may be important in reinforcing the connection between the SCA and torso, potentially through burying the conserved Trp deeper inside the Trp-cage in comparison with Class I and II HelDs (boxed green residues, Figure 4B). Structural modelling also shows the SCA of *M. xanthus* HelD (HelD_MX_) is longer than that of *M. smegmatis* (HelD_MS_), but shorter than the *B. subtilis* protein (HelD_BS_). The tip of the SCA of HelD_MX_ does not reach the active site (catalytic Mg^2+^, green sphere; compare dashed circles in Figure 5C-F) but would clash with the bridge helix in RNAP (teal, Figure 5D and F), potentially causing it to distort and displace the template DNA strand as seen with HelD_BS_ (Newing et al., 2020). The RNAP trigger loop contains a large insertion in the *Deltaproteobacteria* (β’In6, Figure 5B) similar to that seen in *Gammaproteobacteria*, and it was assumed this (and the βIn4 insertion, Figure 5B) would sterically interfere with HelD binding to RNAP in Gram-negative bacteria. Although the trigger loop in the modelled *M. xanthus* RNAP–HelD complex does clash with HelD_MX_ (Figure 5E and F), this is not extensive and given the inherent flexibility in this domain, small conformational changes would readily enable binding as seen in Gram-positive bacteria (Kouba et al., 2020, Newing et al., 2020, Pei et al., 2020). The CA of HelD_MX_ is similar in size to that of HelD_MS_ (although it does not contain a PCh domain; Figure 5B). The CA domain is required for clamp opening and DNA release in the Gram-positive systems, and likely will serve a similar function in Class III HelDs.

**Figure 5.**
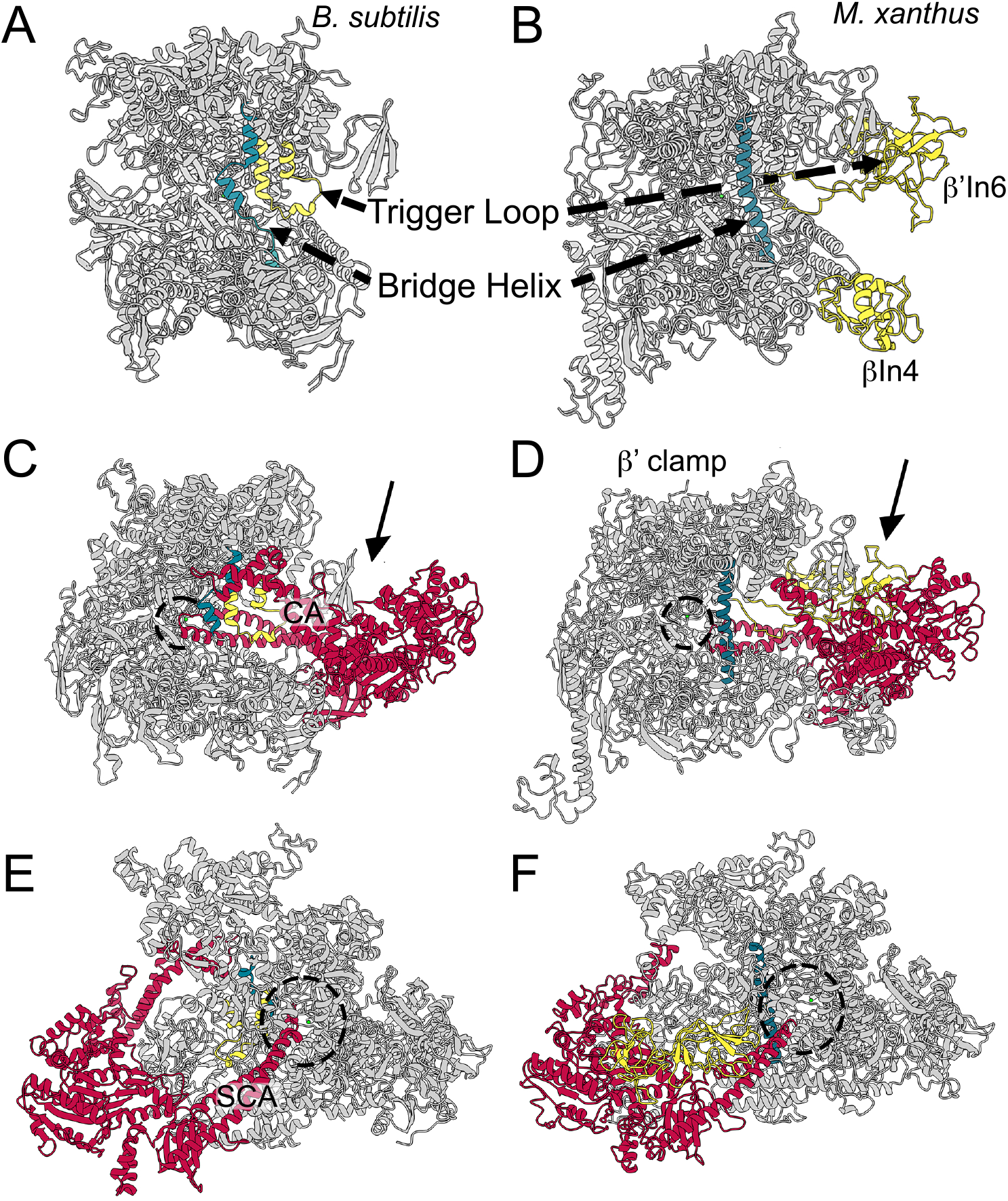
Comparison of *B. subtilis* RNAP–HelD complex with the *M. xanthus* model. Panels A and B show structures of *B. subtilis* (PDB ID 6WVK) and *M. xanthus* (model) RNAPs in complex with HelD, respectively, in which HelD has been removed to more clearly visualise elements referred to in the text. The trigger loop (yellow) and bridge helix (teal) are indicated along with the lineage specific βIn4 (also yellow) and β’In6 inserts in the *M. xanthus* model. Panels C and E show the *B. subtilis* RNAP–HelD complex, PDB ID 6WVK. Panels D and F show *M. xanthus* RNAP–HelD model. RNAP is shown in grey in all panels, HelD in red, bridge helix in teal and trigger loop in yellow (see text for further details). The active site Mg^2+^ is shown as a small green sphere (within the dotted circles). The arrows in panels C and E denote the view of the respective RNAP–HelD complex in panels E and F. The view in panels C and D is into the primary channel to which the clamp arm (CA) of HelD binds. The view in panels E and F is into the secondary channel (dotted circle) into which the secondary channel arm (SCA) is inserted.

Examination of sequences retrieved from the CDART search indicated *helD* genes may be even more widely distributed in the *Proteobacteria* (including the *Gammaproteobacteria*), although this could not be verified by searches of complete genomes in databases such as KEGG and may represent mis-classification from metagenomic sequencing projects. For example, BLASTP searches suggest hits reported as being from *E. coli* and *Vibrio vulnificus* identified from metagenomic data are in fact from *Bacteroides* and *Bacillus*, respectively ((Poyet et al., 2019), and NCBI SRA accession code: PRJNA523266). Nevertheless, it is possible that *helD* genes are more widely distributed in *Proteobacteria*.

### RNAP δ subunit and HelD

The *Firmicutes* have the smallest multi-subunit RNAPs currently known (Lane and Darst, 2010b, Lane and Darst, 2010a), as well as auxiliary subunits δ and ε that are not found in other bacteria (Keller et al., 2014, Weiss and Shaw, 2015). In the original work characterising the function of HelD as a transcription complex recycling factor, it was shown that although δ or HelD on their own enhanced recycling, there was a synergistic relationship between them in *B. subtilis* transcription recycling assays (Wiedermannova et al., 2014). Structural analysis of RNAP recycling complexes shows that δ and HelD interact, as well as providing clues as to how δ could enhance the recycling activity of HelD by augmenting clamp opening (Pei et al., 2020). These structural studies also provided insights into how δ could facilitate transcription recycling in the absence of HelD (Miller et al., 2021). Genome searches indicated that not all *Firmicutes* contained both *helD* and *rpoE* (encoding the δ subunit) genes, and an analysis was performed based on the *rpoB* gene to establish whether there is segregation of genes amongst orders and/or based on natural environment (Figure 6).

**Figure 6.**
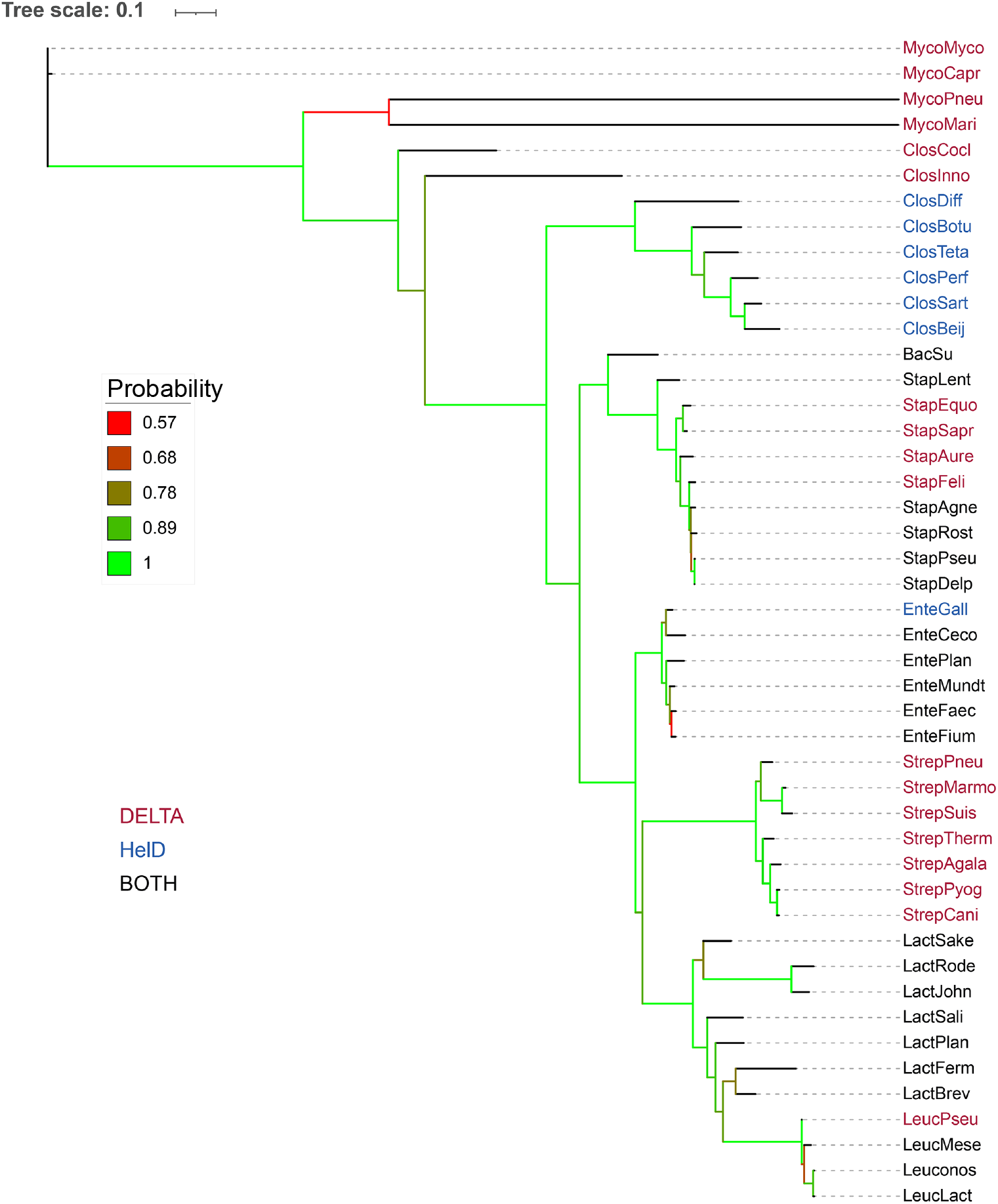
Phylogenetic tree of RpoB with respect to distribution of HelD and the δ subunit of RNAP. Tree scale and bootstrap values are shown on the left. Organisms that contain the δ subunit (DELTA) are shown in red, just HelD (blue) and both δ and HelD (black). *Mycoplasma mycoides* (MycoMyco), *Mycoplasma capricolum* (MycoCapr), *Mycoplasma pneumoniae* (MycoPneu), *Mycoplasma marinum* (MycoMari), *Erysipelatoclostridium cocleatum* (ClosCocl), *Erysipelatoclostridium inoccuum* (ClosInno), *Clostridioides difficile* (ClosDiff), *Clostridium botulinum* (ClosBotu), *Clostridium perfringens* (ClosPerf), *Clostridium sartagoforme* (ClosSart), *Clostridium beijernickii* (ClosBeij), *Bacillus subtilis* (BacSu), *Staphylococcus lentus* (StapLent), *Staphylococcus equorum* (StaphEquo) *Staphylococcus saprophyticus* (StapSapr), *Staphylococcus aureus* (StapAure), *Staphylococcus felis* (StapFeli), *Staphylococcus agnetis* (StapAgne), *Staphylococcus rostri* (Staprost), *Staphylococcus pseudointermidius* (StapPseu), *Staphylococcus delphini* (StapDelp), *Enterococcus gallinarum* (EnteGall), *Enterococcus cecorum* (EnteCeco), *Enterococcus plantarum* (EntePlan), *Enterococcus mundti* (EnteMundt), *Enterococcus faecalis* (EnteFaec), *Enterococcus faecium* (EnteFium), *Streptococcus pneumoniae* (StrepPneu), *Streptococcus marmotae* (StrepMarmo), *Streptococcus suis* (StrepSuis), *Streptococcus thermophilus* (StrepTherm), *Streptococcus agalactiae* (StrepAgala), *Streptococcus pyogenes* (StrepPyog), *Streptococcus canis* (StrepCani), *Lactobacillus sakei* (LactSake), *Lactococcus rodentium* (LactRode), *Lactobacillus johnsonii* (LactJohn), *Lactobacillus salivarius* (LactSali), *Lactobacillus plantarum* (LactPlan), *Lactobacillus fermentum* (LactFerm), *Lactobacillus brevis* (LactBrev), *Leuconostoc pseudomesenteroides* (LeucPseu), *Leuconostoc mesenteroides* (LeucMese), *Leuconostoc sp*. (Leuconos), and *Leuconostoc lactis* (LeucLact).

In the bulk of cases, the *Bacilli, Lactobacilli, Leuconostoc* and *Enterococci* contained genes for both HelD and δ, and if the gene for one protein was missing, the other was present (Figure 6). The *Staphylococci* were heterogeneous with species such as *S. rostri* containing both *helD* and *rpoE* genes, whereas *S. aureus* only contained the gene for the δ subunit. There is a segregation of species containing both *helD* and *rpoE cf. rpoE* only, with *rpoE* only present in the *S. saprophyticus* and *S. aureus* clusters (Takahashi et al., 1999). Species that fall within the *S. hyicus-intermedius* cluster (*e*.*g*., *S. rostri*) contained both *helD* and *rpoE*, but there were exceptions such as *S. felis*, which only contained *rpoE* (Figure 6). The *Streptococci* (order *Lactobacillales*) only contained the *rpoE* gene (Figure 6), whereas the *Clostridia*, except for *C*. (*Erysipelatoclostridium*) *cocleatum* and *inoccuum*, only contained *helD* genes (Figure 6). Thus, it appears that in the *Firmicutes*, especially class *Bacillus*, the default situation is for both *rpoE* and *helD* to be present, but the absence of one gene is compensated for by the presence of the other.

### Many bacteria contain multiple *helD* genes

A striking observation made in the preliminary phylogenetic analysis of HelD was that some organisms contain more than one *helD* gene (Newing et al., 2020). This preliminary analysis has now been extended and it is clear that the presence of >1 *helD* is common and is found in both Gram-positive and -negative organisms (Figure 3A). Using complete genome sequences, up to 5 genes encoding HelD have been identified (*e*.*g. Nonomuraea sp*. ATCC55076 [organism 55]; Figures 3A and S6), and organisms have been identified with 1, 2, 3, 4, or 5 *helD* genes. Although most contain a single *helD* gene, low G+C Gram-positives and Gram-negatives were not found with >3, and high G+C Gram-positive *Actinobacteria* such as *Streptomyces, Nonomuraea*, and *Frankia* were identified with ≥4 *helD* genes. A simple assumption is that these multiple genes are the product of amplification through recombination, and this may well be the root of their original source, but phylogenetic analysis indicates each gene is unique, and organisms with more than one *helD* gene tend to encode both large (∼740-850 aa) and small (∼680-720 aa) variants. The variation in sequence length is due to differences in the flanking SCA and CA domains (arms) with the core 1A and 2A helicase domains all being of similar size. This suggests the motor function of these proteins is conserved, but the function of large *vs* small HelD variants may differ depending on the size of the SCA and CA arms. The multiple *helD* genes also segregate to Class I, -II, or -III according to the organism in which they are found; Class I sequences are found in *Firmicutes*, whereas *Actinobacteria* all have Class II sequences (with the exception of the Coriobacterium *Adlercreutzia equolifaciens*, above), and Class III sequences are found in *Deltaproteobacteria*. Of the *Bacteroides/Parabacteroides* analysed to date, all encode only a single Class I *helD* gene.

It was possible that some/all of the additional *helD* sequences represented cryptic genes that are not expressed under any conditions, or that they are differentially expressed during different growth phases or conditions, which might provide clues to potential functions. Transcriptomics data were retrieved from the Sequence Read Archive (SRA) for selected organisms containing 1 or >1 *helD* representative of all three classes of HelD, and expression levels compared relative to *rpoB* (RNAP β subunit) and another housekeeping gene (SF1 helicase *pcrA/uvrD*). In all cases, all of the *helD* genes were expressed, often at an approximately similar level to *pcrA/uvrD* (Figure 7). The RNA-seq data of *B. subtilis helD* and *pcrA* obtained from experiments by Revilla-Guarinos et al. (Revilla-Guarinos et al., 2020) to examine changes in gene expression in a model soil organism on exposure to the anti-fungal agent amphotericin B produced by *Streptomycetes* closely matched that of the oligonucleotide hybridisation transcriptomics data of Nicolas et al. (Nicolas et al., 2012) and showed the level of *helD* expression was not influenced by amphotericin B and was ∼3% that of *rpoB* (Figure 7A). This is also consistent with proteomics analysis indicating HelD is present at ∼6% the level of RNAP (Delumeau et al., 2011). *B. cereus* contains two *helD* genes and the data set from strain F837/76 (Jessberger et al., 2019) grown in the presence and absence of mucin that can influence toxin production shows that both copies (one large, one small variant) are expressed, albeit at low levels, and expression is not significantly affected on exposure to mucin (Figure 7B). *C. perfringens* also contains two Class I *helD* genes, labelled CPE_0599 (small; 706 aa) and CPE_1619 (large; 763 aa) in strain 13, and expression levels were determined from datasets of cells grown in brain heart infusion (BHI) and a rich medium developed for the optimal growth of fastidious anaerobes, fastidious anaerobe broth + 2% glucose (FABG) medium (Soncini et al., 2020). Both genes were expressed at levels comparable to *helD* in *B. subtilis*, and their cognate *prcA/uvrD*, although CPE_0599 expression increased ∼3-fold and CPE_1619 expression decreased in FABG medium compared to BHI medium (Figure 6C).

**Figure 7.**
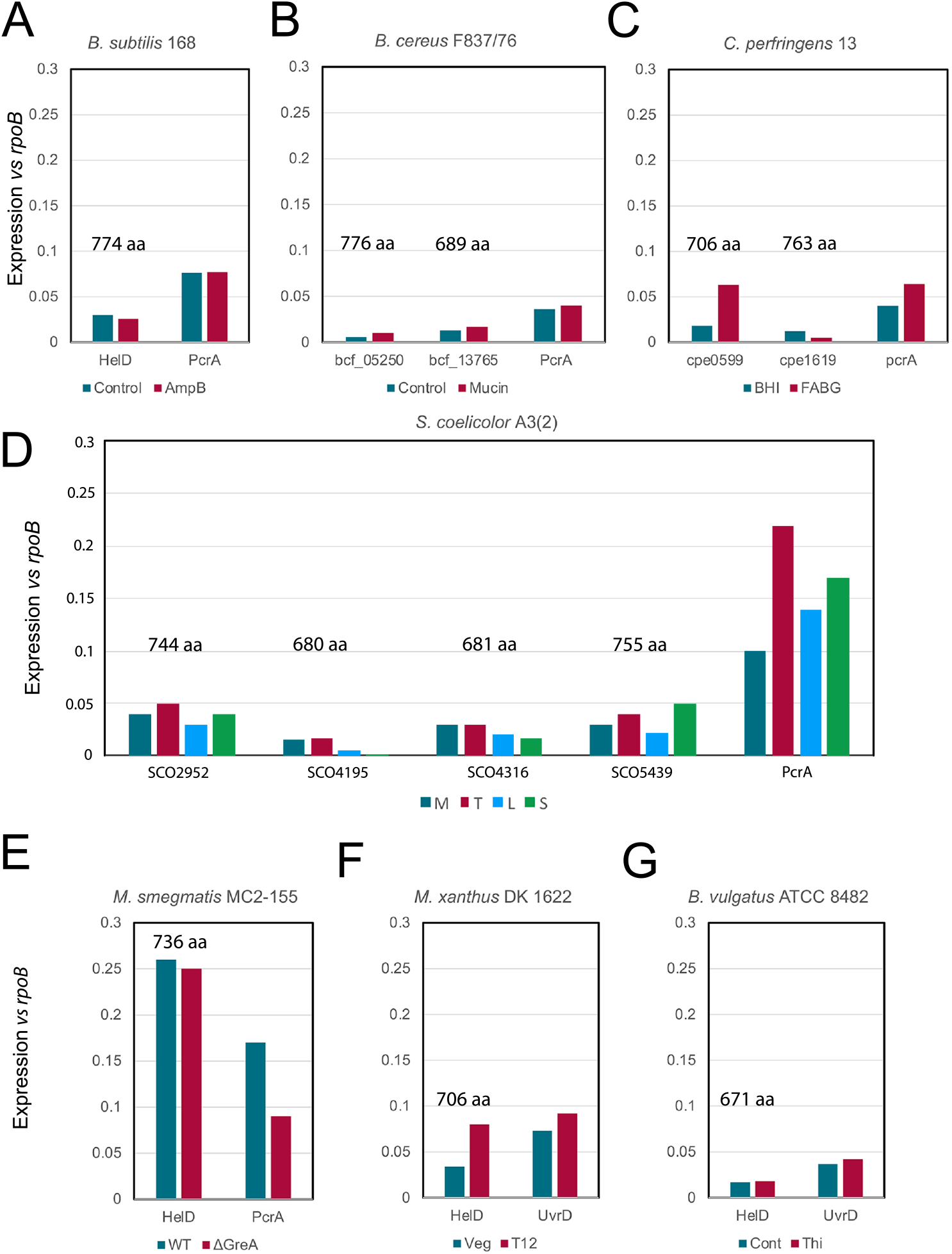
Expression levels of HelD. The relative transcript levels of *helD* and *pcrA/uvrD* compared to *rpoB* are shown in panels A–G. Organism names are shown on the top of each plot and gene expression levels are colour coded according to the keys below the plots. The sizes of the HelD isoforms in amino acids are indicated above the corresponding column in each panel. Details of the sources of the data sets used are provided in the text. A. *B. subtilis* 168 data; control teal, amphotericin B (AmpB) treatment red. B. *B. cereus* F837/76 data; control teal, mucin treatment red. C. *C. perfringens* 13 data; growth in brain heart infusion (BHI) teal, fastidious anaerobic broth + glucose (FABG) red. D. *S. coelicolor* A3(2) data; mid-exponential growth (M) teal, transition phase (T) red, late exponential (L) blue, stationary phase (S) green. E. *M. smegmatis* MC2-155 data; control teal, *greA* deletion strain (ΔGreA) red. F. *M. xanthus* DK1622 data; vegetative growth teal, 12 hours after initiation of sporulation (T12) red. G. *B. vulgatus* ATCC 8482 data; control teal, supplemented with thiamine (Thi) red.

*S. coelicolor* A2(3) contains four Class II *helD* genes, two encoding large (SCO_2952 744 aa, and SCO_5439 755 aa) and two encoding small (SCO_4195 680 aa, and SCO_4316 681 aa) variants. Data from a study on growth phase-dependent changes in gene expression (Jeong et al., 2016) were obtained from the SRA for analysis of *helD* expression and compared with *rpoB* and *pcrA*. All four *helD* genes were expressed with relative levels changing ∼2-fold dependent on the growth phase (Figure 7D). Expression levels were generally highest during mid-log and transition, and lowest during late and stationary phases, with modest changes between the ratios of expression of the different gene copies at all stages. The RNA-seq data set for *M. smegmatis* comparing changes in gene expression on deletion of the transcript cleavage factor GreA that is important in rescuing back-tracked RNAP (Feng et al., 2020) showed that expression of the single *helD* gene was substantially higher than in most other organisms, at about 25% the level of *rpoB* suggesting HelD may be particularly abundant in the *Mycobacteria* (Figure 7E). The expression levels of *helD* were similar in the presence and absence of *greA* indicating each factor acts on stalled transcription complexes independently of each other.

Analysis of RNA-seq data showed *helD* genes were also expressed in Gram-negative *M. xanthus* and *B. vulgatus* (Figure 7F and G), showing that despite the structural differences adjacent to the HelD interaction sites in the β and β’ subunits of RNAP from these organisms, HelDs are expressed and likely able to bind and functionally interact with their cognate RNAPs. The data for *M. xanthus* were obtained to examine changes in gene expression during the development of fruiting bodies and spores. It is interesting to note that expression of *helD* in *M. xanthus* increases during development of spores (not to be confused with sporulation in the *Firmicutes*) and may point to a role in storage of inactive RNAP during dormancy as has been proposed for *B. subtilis* HelD (Pei et al., 2020). The study in *B. vulgatus* was designed to investigate the effect on gene expression of exogenous thiamine that may be important in niche establishment in the gut. Therefore, in most/all organisms that contain *helD* gene(s), it/they are expressed. The reason why one organism contains a single gene and closely related species contain more than one (*e*.*g. B. subtilis* and *B. cereus*, Figure 6A and B) is currently not clear, but the expression data would suggest that each isoform has a functional role to play in the cell, and there is not a significant difference in the expression of large *vs* small *helD* variants.

## CONCLUSIONS

In this work we have examined the phylogenetic distribution and classification of the transcription recycling factor HelD in detail and have identified a new class restricted to the *Deltaproteobacteria*. In addition, it appears *helD* genes have been acquired by horizontal transfer on at least three occasions; *Bacteroides* have acquired *helD* from the *Clostridiales*, whereas the *Coriobacteria* have acquired it from the *Lactobacilli* and *Clostridiales*. The gut microbiome is known as an environment conducive to horizontal gene transfer, especially with respect to distribution of antibiotic resistance genes (McInnes et al., 2020), and given that *Bacteroides, Lactobacilli, Clostridiales*, and *Coriobacteria* are all common in the gut microbiome, it appears *A. equolifaciens* has aquired *helD* genes from gut microorganisms on two separate occasions. Indeed, an unusual feature of *helD* genes is that many organisms contain multiple paralogues, and that all versions are expressed. Why some organisms have a single gene for *helD* while a closely related species has multiple expressed copies is unclear, and this will make a fascinating avenue for future research. It is interesting to note that actinobacteria, such as *Streptomyces, Frankia*, and *Nonomuraea* (numbers 50, 51, 54 and 55; Figure. 1) that are known producers of valuable bioactive compounds used as antibiotics and anti-cancer drugs contained the largest number of *helD* genes (4-5). It is possible that the 5 *helD* genes in *Nonomuraea* (number 55, Figure 1), that is a known producer of DNA-intercalating agents (Sungthong and Nakaew, 2015) are involved in genome maintenance through recycling stalled transcription complexes during production of these compounds. *Nonomuraea* and other *Actinomycetales* sometimes have a second *rpoB* gene that confers resistance of RNAP to compounds such as rifampicin and sorangicin that is induced by stress and is associated with production of secondary metabolites (D’Argenio et al., 2016). The combination of multiple HelD isoforms with drug resistant RNAP may be important in this proposed genome maintenance activity. In some organisms, such as *M. abcessus* and *S. venezuaelae helD* expression is induced in the presence of the antibiotic rifampicin, conferring resistance, and this is associated with the presence of a DNA sequence called the Rifamycin Associated Element (RAE) found upstream of the gene (Hurst-Hess et al., 2021, Surette et al., 2021). It is proposed that the tip of the PCh loop is able to physically remove rifampicin bound to the RNAP β subunit in a pocket close to the active site. In *S. venezuelae* (organism #50, Figure 3) that has five *helD* genes, only one (SVEN_6029, #50.3) is induced in the presence of rifampicin and has an upstream RAE (Surette et al., 2021). It is interesting to note that despite encoding a rifampicin resistant RNAP β subunit, *Nonomuraea* also has an RAE located directly upstream of *helD* NOA_42280 (#55.3; Figure S5).

Investigation of the distribution of *helD* genes with upstream RAEs revealed they were clustered to two sub-branches of the *Actinobacteria* (Figure S7) that may be considered the HelR grouping based on the nomenclature of these proteins by (Hurst-Hess et al., 2021, Surette et al., 2021). It should be noted that clearly identifiable RAEs could not be found upstream of all the genes in the HelR group, including for *Frankia alni, Nocardia brasiliensis* or *Mycolicibacterium phlei* (54.2, 56.2, and 64, respectively; Figure 3 and S2). Rifampicin has also been observed to induce *helD* expression in the low G+C Gram-positive *B. subtilis*, but this induction does not confer resistance to the drug (Hutter et al., 2004). Nevertheless, the ability of naturally produced antibiotics to induce expression of *helD* genes suggests HelD proteins have a potentially important role in preserving genome integrity and gene expression in the bacteria in which they are found.

An additional area of future research should include functional and structural studies of HelD from Gram-negative bacteria, as due to the location of lineage-specific inserts in the β and β’ subunits of RNAP in Gram-negatives it was assumed HelD-like proteins would bind poorly or be sterically inhibited from binding. HelD proteins represent a new class of motor enzyme involved in transcription complex recycling that are widely distributed in bacteria that make an important contribution to our understanding of the multiple different mechanisms used to resolve potentially lethal stalled transcription complexes.

Finally, it is important that genome annotation databases are updated as *helD* genes are often classified as *pcrA, uvrD*, or helicase IV-ATPase. Correct annotation of *helD* genes will enable more detailed understanding of the distribution, evolution and function of this fascinating new category of transcription factor.

## EXPERIMENTAL PROCEDURES

### Sequence retrieval and analysis

The sequence of *B. subtilis* 168 HelD (UniProtKB ID: O32215) was used to search for homologues using the NCBI Conserved Domain Architecture Retrieval Tool (Geer et al., 2002), which identified 13,781 sequences, which were trimmed to 11,821. To aid subsequent analyses, particularly for the study of multiple copies of *helD* genes, the original sequences were used to search complete reference genomes from the KEGG (https://www.kegg.jp) and JGI (https://jgi.doe.gov) databases. HelD and RpoB sequences retrieved from these complete genomes were used for subsequent phylogenetic studies.

### Construction of phylogenetic trees

Selected sequences were aligned using MAFFT (Katoh et al., 2002, Katoh et al., 2019) with default settings. Sequence alignments were then trimmed using Gblocks (https://ngphylogeny.fr). The best fitting model (LG) was determined using ProtTest 3 (Darriba et al., 2011) and phylogenetic trees were constructed using MrBayes 3.2 (Huelsenbeck and Ronquist, 2001, Ronquist et al., 2012), which were run until the standard deviation was below 0.01. Trees were visualised using iTol (Letunic and Bork, 2019).

### Transcriptome data and analysis

Gene expression data were obtained from datasets deposited in the Sequence Read Archive (SRA; https://www.ncbi.nlm.nih.gov/sra) and were: *B. subtilis* 168 (Revilla-Guarinos et al., 2020); *B. cereus* F837/76 (Jessberger et al., 2019); *Clostridium perfringens* 13 (Soncini et al., 2020); *Streptomyces coelicolor* A3(2) (Jeong et al., 2016); *Mycobacterium smegmatis* MC2-155 (Feng et al., 2020); *Myxococcus xanthus* DK1622 (SRA accession code: PRJNA516475); *Bacteroides vulgatus* ATCC8482 (SRA accession code: PRJNA473003). Reads were mapped to the respective reference genome sequences, and gene expression levels were calculated in Genious Prime 2020.2.3 (https://www.geneious.com). Transcript per million (TPM) values were used for comparison of *helD* expression levels *cf. rpoB*, and *pcrA/uvrD* (for *S. coelicolor* A3(2)).

### Structure modelling

RNAP RpoB (β) and RpoC (β’) subunits from *M. xanthus* DK1622 were modelled in SWISS-MODEL (Waterhouse et al., 2018) using *E. coli* RNAP, PDB ID: 6ALF (Kang et al., 2017) as a defined template. The *M. xanthus* HelD structure was modelled using i-Tasser (Yang et al., 2015) with output model 1 (C-score -0.48) selected for presentation in this work. Structural images used in this work were prepared in ChimeraX (Pettersen et al., 2020).

## ACKNOWLEDGEMENTS

The authors appreciate the constructive comments from Brett Neilan, Leanne Pearson-Neilan, Caitlin Romanis and Karl Hassan during the preparation of this manuscript.

## AUTHOR CONTRIBUTIONS

JSL and MM, acquisition, analysis and interpretation of data. AJO, analysis and interpretation of data. NED, interpretation of data and writing of manuscript. PJL, conception and design of study, acquisition, analysis and interpretation of data, writing of manuscript.

## DATA AVAILABILITY

The hybrid *M. xanthus* RNAP-HelD complex model is available on request from P.J.L.

## SUPPORTING INFORMATION

Supporting information is available online.

