## Supplementary Material for "Multiple classes and isoforms of the RNA polymerase recycling motor protein HelD"

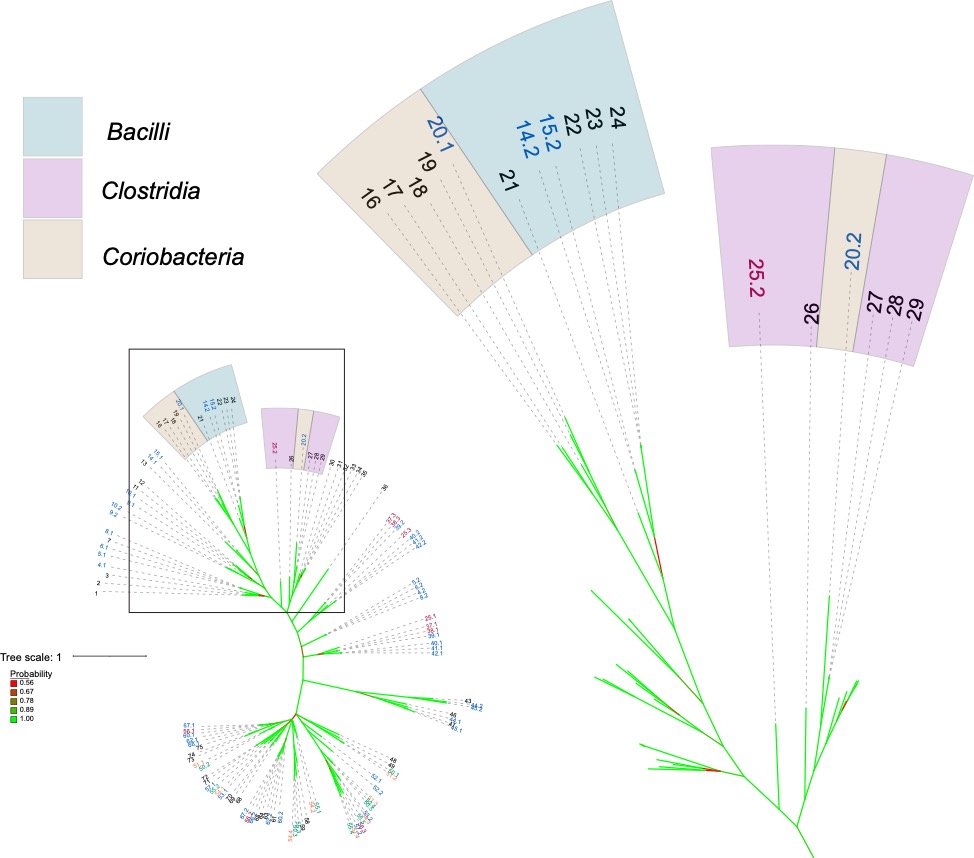

**Figure S1.** Acquisition of *helD* genes by *Coriobacteria* from *Firmicutes* and *Clostridia*. The phylogenetic tree from Figure 1 is shown on the left side with the region boxed expanded on the right side. Bacterial classes are coloured, species numbered, number of *helD* genes coloured as in Figure 1: *Bacilli*, teal; *Clostridia*, purple; *Coriobacteria*, brown. **14** *Enterococcus faecium* Aus0004 (#2 EFAU004_00387, 711 aa). **15** *Enterococcus faecium* DO (#2 HMPREF0351_10397, 711 aa). **16** *Olsenella uli* DSM 7084 (OLS_0501, 731aa). **17** *Atopobium parvulum* DSM 20469 (Apar_0360, 736aa). **18** *Slackia heliotrinireducens* DSM 20476: (Shel_05840 (698aa). **19** *Eggerthella lenta* DSM 2243(Elen_2835, 716aa). **20** *Adlercreutzia equolifaciens* DSM 19450 (#1 AEQU_1689, 761aa; #2 AEQU_0484, 733aa). **21** *Vagococcus teuberi* (vte_03205, 717aa). **22** *Enterococcus faecalis* V583 (EF_0933, 732 aa). **23** *Enterococcus faecalis* DENG1 (DENG_00988, 732 aa). **24** *Enterococcus faecalis* OG1RF (OG1RF_10660, 740 aa). **25** *Clostridium beijerinckii* NCIMB 8052 (#2 cbe_2724, 745aa). **26** *Epulopiscium sp*. N.t. morphotype B (EPU_RS03295, 735aa). **27** *Clostridioides difficile* 630 (CD630_04550, 704 aa). **28** *Clostridioides difficile* RM20291 (CDR20291_0396, 704 aa). **29** *Clostridioides difficile* CD196 (CD196_0410, 704 aa). One *helD* gene, black; two, blue; three, red.

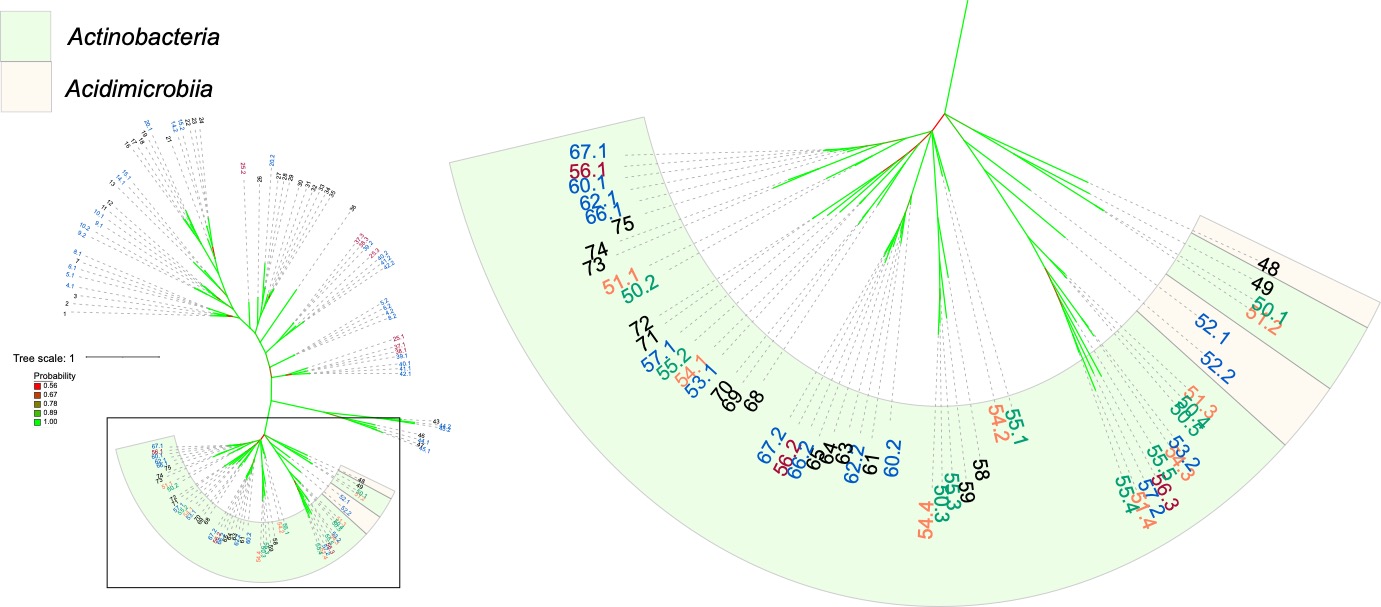

**Figure S2.** Acquisition of *helD* genes by *Acidimicrobiia* from *Actinobacteria*. The phylogenetic tree from Figure 1 is shown on the left side with the region boxed expanded on the right side. Bacterial classes are coloured, species numbered, number of *helD* genes coloured as in Figure 1: *Actinobacteria*, pale green; *Acidimicrobiia*, pale yellow. **48** *Acidobacterium ferrooxidans* (Afer_1829, 706aa). **49** *Cutibacterium acnes* KPA171202 (PPA0733, 753aa). **50** *Streptomyces venezuelae* (#1 SVEN_2719, 779aa; #2 SVEN_5092, 747aa; #3 SVEN_6029, 722aa; #4 SVEN_4127, 675aa; #5 SVEN_3939; 665aa). **51** *Streptomyces coelicolor* A3(2) (#1 SCO5439, 755 aa; #2 SCO2952, 744 aa; #3 SCO4316, 681 aa; #4 SCO4195, 680 aa). **52** *Ilumatobacter coccineus* (#1 aym_09360, 715aa; #2 aym_20540, 654aa). **53** *Frankia casuarinae* Ccl3 (#1 fra_0952, 829aa; #2 fra_2397, 727aa). **54** *Frankia alni* ACN14a (#1 fal_1589, 939aa; #2 fal_4723, 877aa; #3 fal_3805; 866aa; #4 fal_4811, 751aa). **55** *Nonomuraea sp.* ATCC55076 (#1 NOA_23645, 772 aa; #2 NOA_16240, 762 aa; #3 NOA_42280, 715 aa; #4 NOA_08745, 660 aa; #5 NOA_48960, 655 aa). **56** *Nocardia brasiliensis* O31_020410 (#1 nbr_012985, 776aa; #2 nbr_020410, 731aa; #3 nbr: O3I_005870, 699aa). **57** *Kineococcus radiotolerans* SRS30216 (#1 kra_3607, 759aa; #2 kra_0164, 684aa). **58** *Microbacterium sp.* PAMC 28756 (mip_00070, 717aa). **59** *Mirobacterium hominis* SJTG1 (mhos_01135, 744aa). **60** *Nocardia farcinica* IFM10152 (#1 NFA_19060, 765aa; #2 NFA_44160, 726aa). **61** *Mycobacterium smegmatis* MC2 155 (MSMEG_2174, 736aa). **62** *Rhodococcus sp.* 008 (#1 rhod_26990, 760aa; #2 rhod_09075, 731aa). **63** *Mycobacterium sp.* JS623 (Mycsm_03949, 732aa). **64** *Mycolicibacterium phlei* (MPHL_03003, 726aa). **65** *Mycobacteroides abscessus* ATCC 19977 (MAB_3189c, 753aa). **66** *Rhodococcus equi* 103S (#1 REQ_25070, 759aa; #2 REQ_15310, 739aa). **67** *Nocardia asteroides* NCTC11293 (#1 nad_03000, 753; #2 nad_04408, 735aa). **68** *Leifsonia xyxli* subsp. Xyli CTCB07 (Lxx_20770, 787aa). **69** *Bifidobacterium longum* NCC2705 (BLO_1314, 759aa). **70** *Bifodobacterium bifidum* PRL2010 (bbp_0546, 759aa). **71** *Brevibacterium linens* BS258 (bly_10570, 743aa). **72** *Brevibacterium flavum* ZL-1 (bfv_07580, 755aa). **73** *Corynebacterium glutamicum* ATCC13031 (CG_1555, 755aa). **74** *Corynebacterium diptheriae* NTCC13129 (DIP_1156, 770aa). **75** *Rhodococcus rhodochrous* NCTC10210 (rrt_02795, 772aa). One *helD* gene, black; two, blue; three, red; four, orange; five, green.

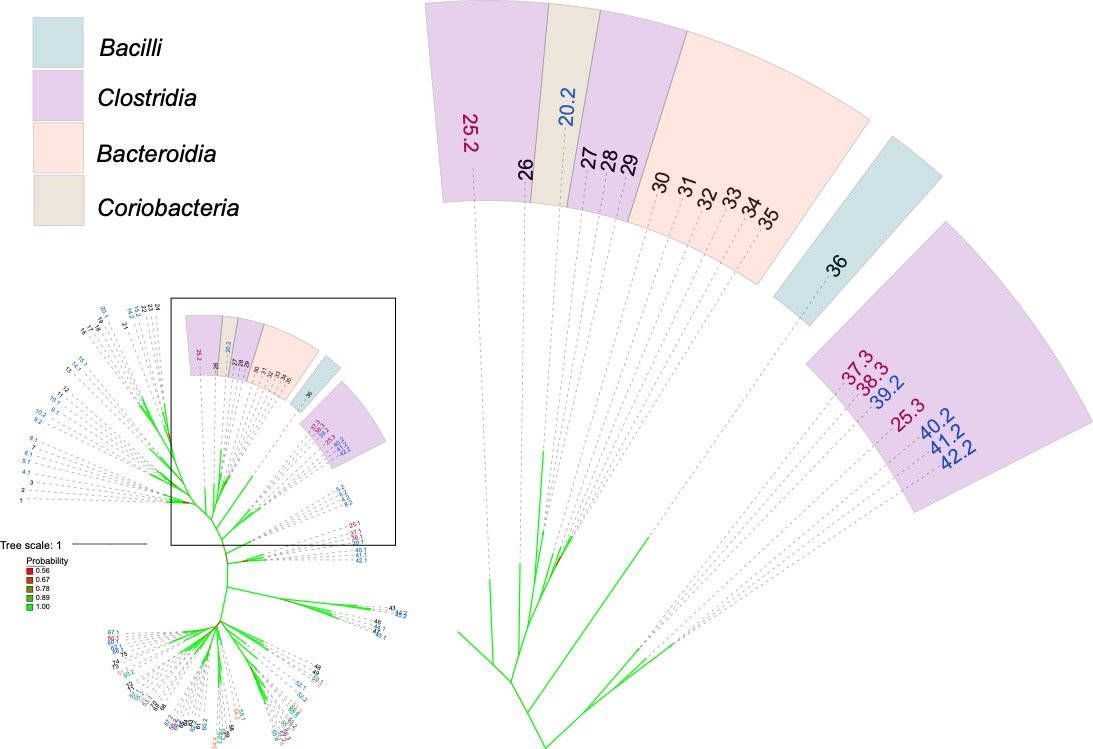

**Figure S3.** Acquisition of *helD* genes by *Bacteroides* from *Clostridia*. The phylogenetic tree from Figure 1 is shown on the left side with the region boxed expanded on the right side. Bacterial classes are coloured, species numbered, number of *helD* genes coloured as in Figure 1: *Bacilli*, teal; *Clostridia*, purple; *Bacteroides*, orange; *Coriobacteria*, brown. **20** *Adlercreutzia equolifaciens* DSM 19450 (#2 AEQU_0484, 733aa). **25** *Clostridium beijerinckii* NCIMB 8052 (#2 cbe_2724, 745aa; #3 cbe_4782, 724aa). **26** *Epulopiscium sp*. N.t. morphotype B (EPU_RS03295, 735aa). **27** *Clostridioides difficile* 630 (CD630_04550, 704 aa). **28** *Clostridioides difficile* RM20291 (CDR20291_0396, 704 aa). **29** *Clostridioides difficile* CD196 (CD196_0410, 704 aa). **30** *Bacteroides vulgatus* ATCC 8482 (BVU_3010 (671aa). **31** *Bacteroides caccae* ATCC 43185 (CGC64_00555, 683aa). **32** *Bacteroides cellulosilyticus* WH2 (BcelWH2_01491, 693aa). **33** *Bacteroides thetaiotaomicron* VPI-5482 (BT_1890, 686aa). **34** *Bacteroides ovatus* ATCC 8483 (Bovatus_02598 (687aa). **35** *Bacteroides xylanisolvens* XB1A (BXY_17560, 687aa). **36** *Staphylococcus delphini* NCTC12225 (sdp_01978, 681aa). **37** *Clostridium botulinuim A* ATCC3502 (#3 CBO_3341, 709 aa). **38** *Clostridium botulinuim A* ATCC19377 (#3 CLB_3399, 709 aa). **39** *Clostridium botulinuim B1 Okra* (#2 CLD_1179, 709 aa). **40** *Clostridium perfringens* 13 (#2 CPE_0599, 706 aa). **41** *Clostridium perfringens* ATCC13124 (#2 CPF_0580, 706 aa). **42** *Clostridium perfringens* SM101 (#2 CPR_0566 706 aa). One *helD* gene, black; two, blue; three, red.

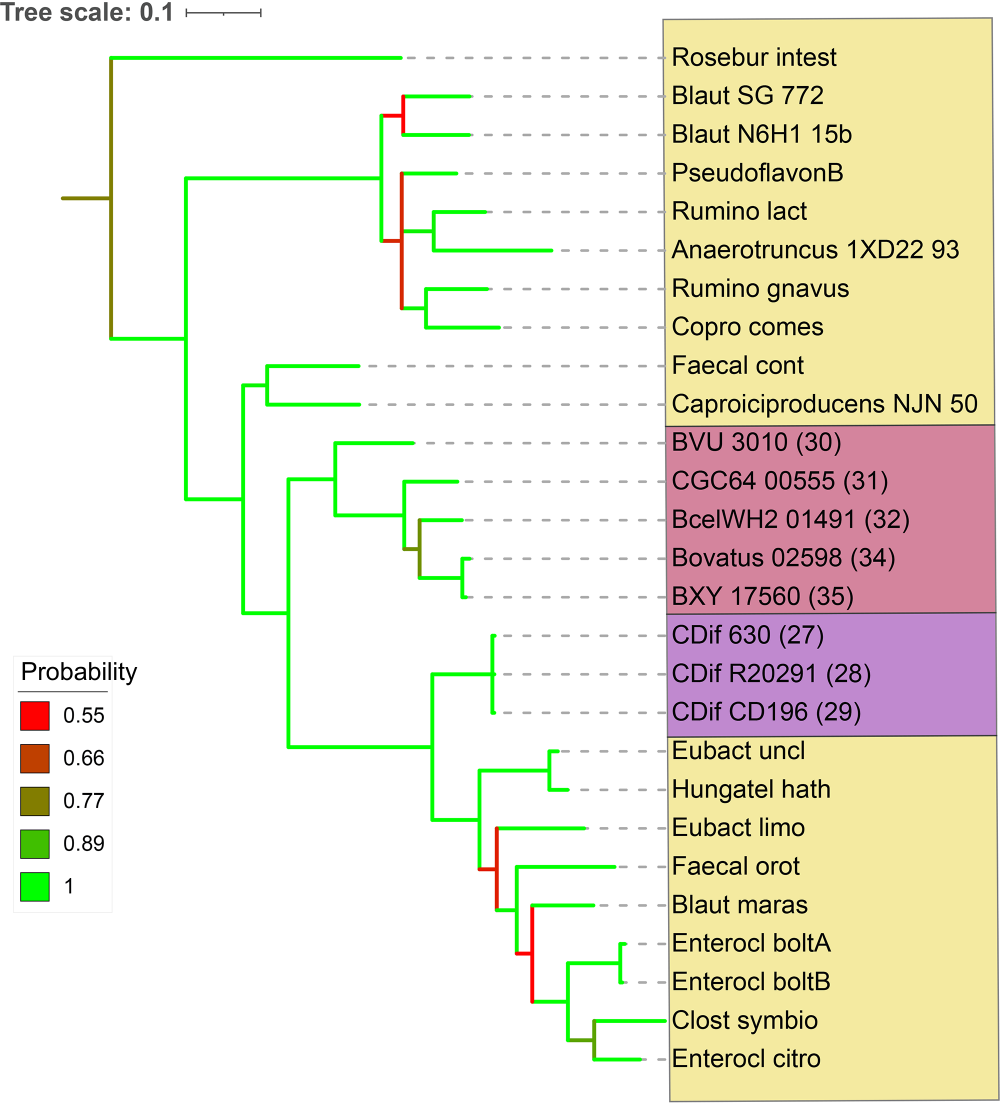

**Figure S4.** Phylogenetic tree of HelD sequences from *Bacteroides* and *Clostridia*. Tree scale and bootstrap values are shown at the top and left, respectively. Coloured boxes denote cluster IV and XIVa *Clostridia* (yellow), cluster IX *Clostridia* (purple), and *Bacteroides* (red). The numbers in parentheses correspond to the organisms used in Figure 1. *Roseburia intestinalis* (Rosebur intest), *Blautia sp.* SG-772 (Blaut SG772), *Blautia sp.* N6H1-15 (Blaut N6H1-15b), *Pseudoflavonifractor sp.* BSD2780061688st1 E11 (PseudoflavonB), *Ruminococcus lactaris* (Rumino lact), *Anaerotruncus sp.* 1XD22-93 (Anaerotruncus 1XD22-93), *Ruminococcus gnavus* (Rumino gnavus), *Coprococcus comes* (Copro comes), *B. vulgatus* ATCC 8482 (BVU 3010), *B. caccae* ATCC 43185 (CGC64 00555), *B. cellulosilyticus* WH2 (BcelWH2 01491), *B. ovatus* ATCC 8483 (Bovatus 02598), *B. xylanisolvens* XB1A (BXY 17560), *C. difficile* 630 (CDif 630), *C. difficile* RM20291 (CDif R20291), *C. difficile* CD196 (CDif CD196), *Faecalicatena contorta* (Faecal cont), *Caproiciproducens sp.* NJN-50 (*Caproiciproducens* NJN-50), *Eubacterium uniforme* (Eubact uncl), *Hungatella hathewayi* (Hungatel hath), *Eubacterium limosum* (Eubact limo), *Faecalicatena orotica* (Faecal orot), *Blautia marasmi* (Blaut maras), *Enterocloster bolteae* (Enterocl boltA and B), *Clostridium symbiosum* (Clost symbio), and *Enterocloster citroniae* (Enterocl citro).

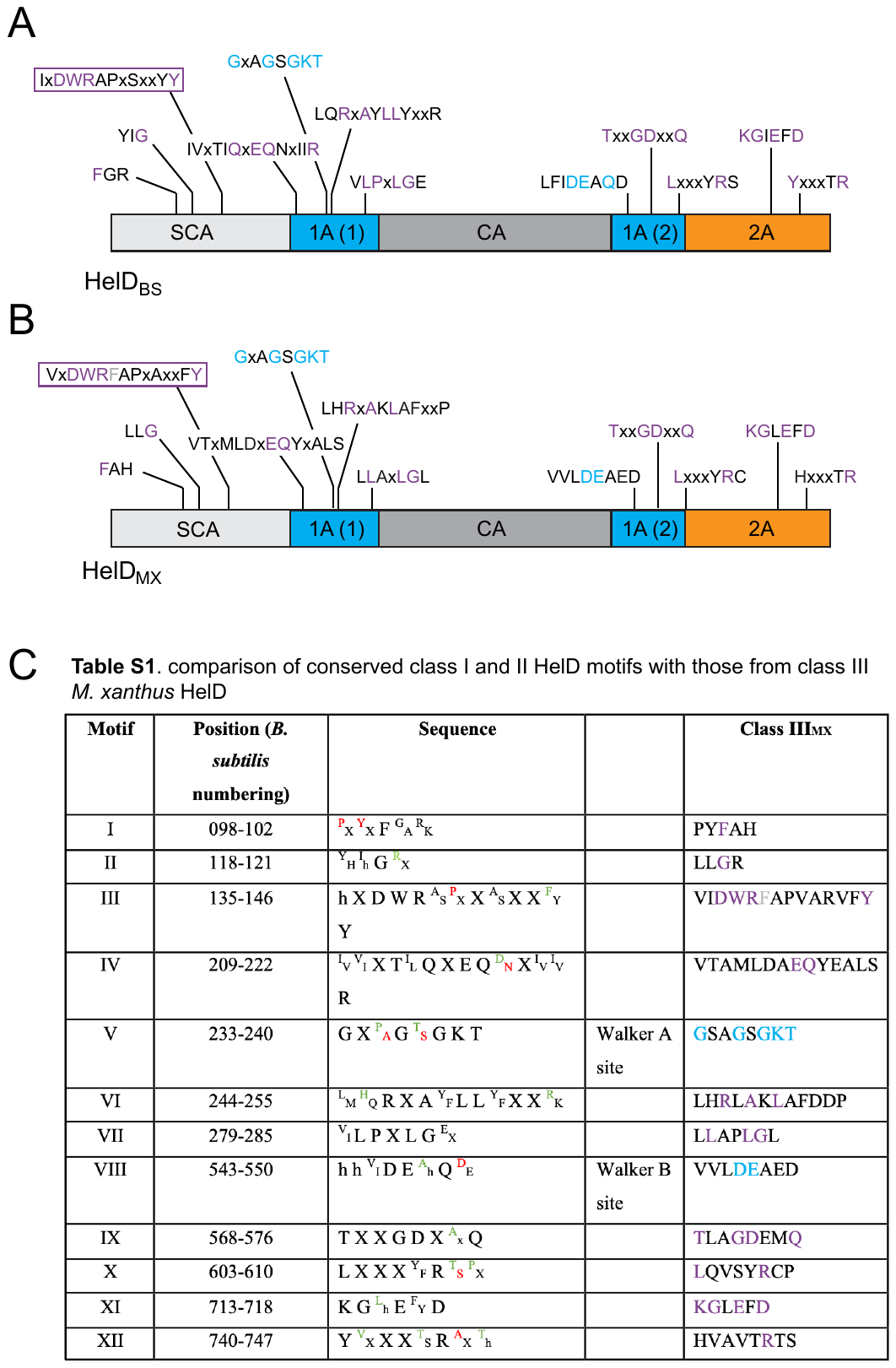

**Figure S5.** Conserved HelD sequence motifs. Panel A shows a schematic of *B. subtilis* HelD domain organisation with conserved sequence motifs adapted from Newing *et al*., {Newing, 2020 #813}, with panel B showing the equivalent sequence motifs from *M. xanthus* HelD. Table S1 shows the conserved sequence motifs with sequence numbers referring to the *B. subtilis* HelD sequence. X corresponds to a poorly conserved sequence (any amino acid) and h to a conserved hydrophobic residue. Residues coloured red are specific to class I and green to class II sequences. The HelD motifs from the Class III *M. xanthus* HelD (Class III_MX_) are shown in the right column with absolutely conserved motif residues shown in purple (blue for the ATP binding motifs) and the Class III defining residue (F in the case of *M. xanthus*) that is inserted in the DWRAP motif shown in grey (see text for more details).

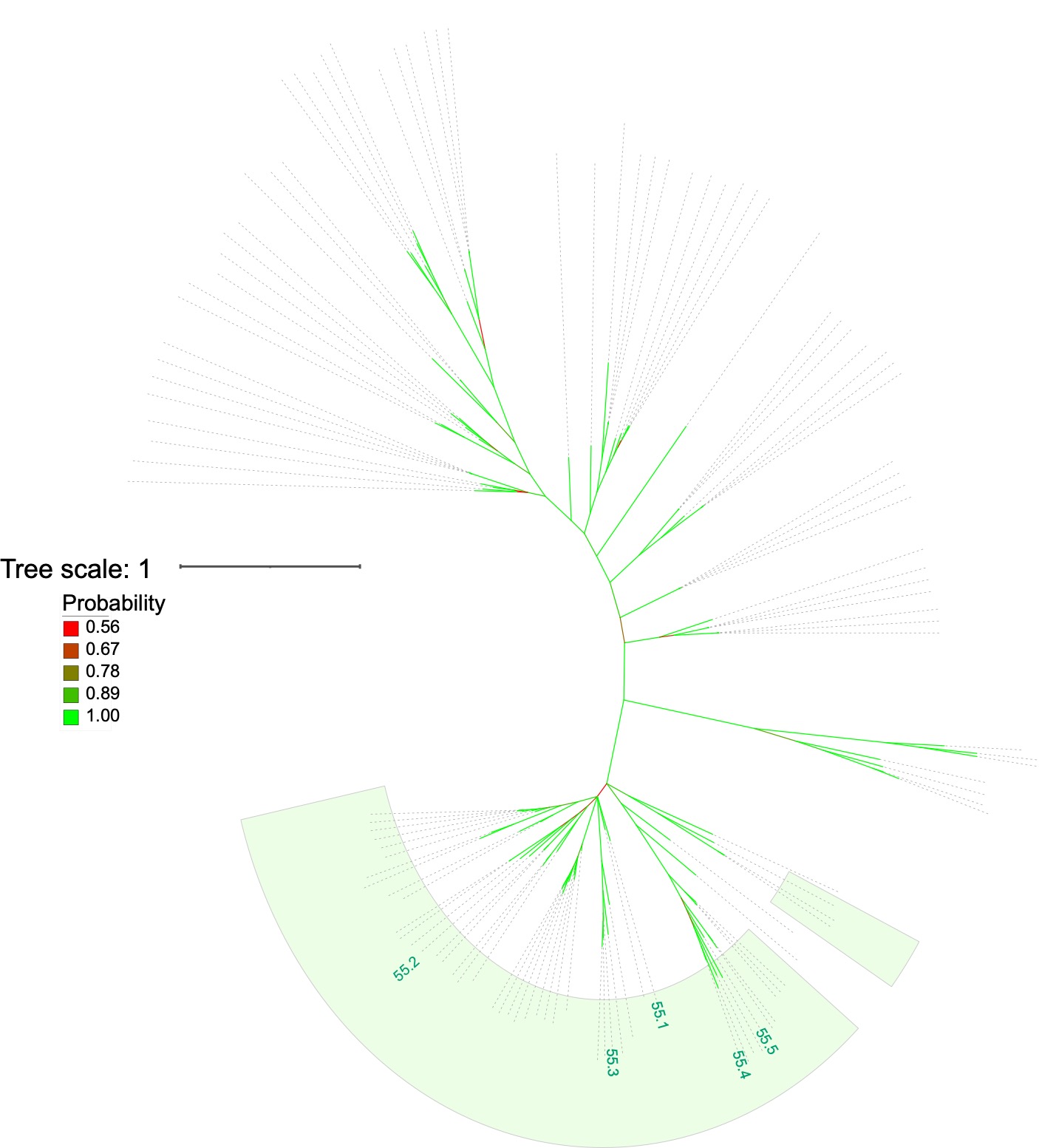

**Figure S6.** Distribution of the five *helD* genes from *Nonomuraea sp.* ATCC55076. The phylogenetic tree from Figure 1 is shown unannotated apart from boxing the region corresponding to the *Actinobacteria* pale green, and indicating the location of the *Nonomuraea helD* genes: #1 NOA_23645, 772 aa; #2 NOA_16240, 762 aa; #3 NOA_42280, 715 aa; #4 NOA_08745, 660 aa; #5 NOA_48960, 655 aa.

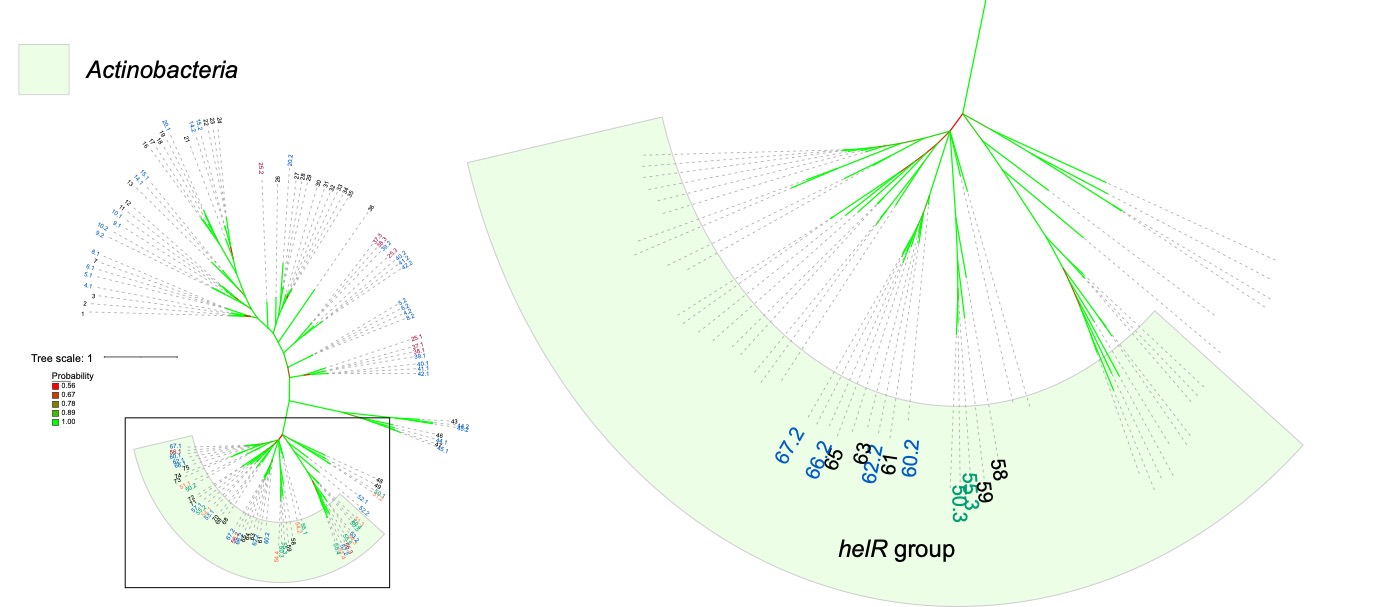

**Figure S7.** The *helR* group of *helD* variants that (potentially) confer rifampicin resistance. The phylogenetic tree from Figure 1 is shown on the left side with the region boxed expanded on the right side. Bacterial classes are coloured, species numbered, number of *helD* genes coloured as in Figure 1. Only the (potential) *helR* variants are shown: **50** *Streptomyces venezuelae* (#3 SVEN_6029). **55** *Nonomuraea sp.* ATCC55076 (#3 NOA_42280, 715 aa). **58** *Microbacterium sp.* PAMC 28756 (mip_00070, 717aa). **59** *Mirobacterium hominis* SJTG1 (mhos_01135, 744aa). **60** *Nocardia farcinica* IFM10152 (#2 NFA_44160, 726aa). **61** *Mycobacterium smegmatis* MC2 155 (MSMEG_2174, 736aa). **62** *Rhodococcus sp.* 008 (#2 rhod_09075, 731aa). **63** *Mycobacterium sp.* JS623 (Mycsm_03949, 732aa). **65** *Mycobacteroides abscessus* ATCC 19977 (MAB_3189c, 753aa). **66** *Rhodococcus equi* 103S (#2 REQ_15310, 739aa). **67** *Nocardia asteroides* NCTC11293 (#2 nad_04408, 735aa). One *helD* gene, black; two, blue; five, green.
